# Large-Vessel Venous Signal Confounds Apparent Amygdala Activation in BOLD fMRI

**DOI:** 10.64898/2026.09.04.749520

**Authors:** Zachary Diamandis, J. Michael Tyszka, Ralph Adolphs

## Abstract

Amygdala responses are widely reported in BOLD fMRI studies of human emotions and mood disorders. Although modern acquisition methods have improved image quality, the hemodynamic nature of the BOLD signal continues to complicate interpretation of these responses. Across an array of both visual and auditory tasks, we here show that apparent amygdala BOLD signal is substantially confounded by signal from nearby deep veins, including the basal vein of Rosenthal. In contrast to prior work, we show that the amygdala, when engaged, is the leading but not exclusive contributor to this venous signal. Across the tested conditions, this signal was sensitive to, but not specific for, amygdala engagement. This confound mislocalizes apparent activation and prevents reliable attribution of signal to amygdala subnuclei with conventional BOLD fMRI. We further show that spatial smoothing exacerbates this mislocalization by merging peri-amygdalar venous signal with the anatomical amygdala. We reproduce our findings from precision imaging in a larger sample obtained from the Human Connectome Project. While broad conclusions about amygdala engagement remain largely valid when based on unsmoothed signal within the anatomical amygdala, further conclusions about subnuclear function drawn from conventional fMRI alone warrant considerable caution.

## Introduction

The amygdala is a complex subcortical region comprising over a dozen distinct nuclei in primates, and has emerged as a central structure in emotional and social cognition. In humans, the amygdala has been the target of a large literature covering a wide array of cognitive processes [1, 2, 3] and psychiatric disorders [4], with functional magnetic resonance imaging (fMRI) being by far the most commonly used approach [1]. This approach has enabled non-invasive study of amygdala responses at scale, in population cohorts of up to tens of thousands of participants [5, 6, 7, 8] and in a wide range of psychiatric populations [4, 9]. The large human neuroimaging literature has supported inferences regarding amygdala function, including increasingly more spatially precise claims at the subnuclear level [10, 11].

Yet amygdala fMRI findings remain inconsistent and difficult to interpret. Studies differ significantly in reported findings – not only in fine-grained spatial detail, but also in test–retest reliability and even whether amygdala activation is reported at all. Recent work has raised broad concerns about the reliability and validity of amygdala fMRI, especially when supporting individual difference or clinical biomarker claims [12, 13, 14]. Other work has shown that lateralization and activation patterns can change with image preprocessing choices, including smoothing and motion correction [15]. In fact, physiological-noise correction has not consistently improved the test–retest reliability of amygdala BOLD responses, with effects varying across tasks and reliability measures [16, 17]. Studies attempting to localize responses within amygdala subregions report strikingly different results. Emotional-versus-neutral face effects have been assigned to the left basolateral and superficial groups [18]; responses to fearful, happy, and neutral faces versus houses to the superficial group bilaterally [19]; face-versus-blank-baseline responses to the accessory basal nucleus bilaterally and an expression-by-context interaction to the right corticomedial group [20]; and emotional-face-versus-shape responses to all four assessed subregions, with the superficial response greater than all other subregions and the centromedial response greater than the basolateral and amygdalostriatal responses [21]. High-resolution fMRI with negative visual stimuli likewise reported significant negative-versus-neutral responses in both the centromedial and basal groups, with the largest response in the centromedial group [22]. Although methodological differences mean that these studies should not be read as direct contradictions, the range of implicated subregions illustrates the heterogeneity of this literature. As a result, much of our validation of amygdala function depends on electrophysiological experiments in rodents and non-human primates. While these findings do not suggest that amygdala fMRI is uninformative, they do warrant caution for fine-grained inference, particularly in terms of anatomical and functional specificity.

Several factors may contribute to the heterogeneity reported from BOLD fMRI studies of the amygdala. The amygdala itself is a small, deep structure within the brain, close to bone- and air-tissue interfaces and far from receiver coil elements. High-quality MRI of the amygdala is intrinsically more challenging than in superficial cortex due to poorer *B*_0_ homogeneity, lower signal-to-noise ratio (SNR), and greater noise amplification from image acceleration [23, 24]. Equally concerning is the amygdala’s proximity to large draining veins such as the deep middle cerebral vein and basal vein of Rosenthal (BVR), with task-modulated BOLD signal from the amygdala potentially reaching the BVR via the amygdalar, inferior ventricular, and uncal veins [25, 26].

BOLD fMRI is an indirect measure of neuronal activity, inherently limited by the reliability of neurovascular coupling to accurately represent underlying neural dynamics [27]. The impact of macrovascular components on the spatial specificity of cerebral BOLD responses has been recognized since the earliest days of BOLD fMRI [28, 29, 30]. A variety of methods for reducing macrovascular bias in BOLD fMRI responses have been proposed subsequently, including moving to higher magnetic field strengths (greater than 3 T) [31], the use of T2-prepared gradient-echo imaging at high field (7 T) [32], and post-processing approaches such as phase regression of complex-valued MRI data [33]. However, the vast majority of published BOLD fMRI studies with subcortical targets do not address macrovascular bias directly with such pulse sequences or post-processing approaches or with higher-field systems.

This issue of large vein contributions to apparent amygdala BOLD responses was first raised by Boubela et al. in 2015, who found that a commonly observed response to an emotional face task was located dorsomedial to the amygdala and was largely consistent with the course of the basal vein of Rosenthal, arguing that such signal could reflect venous drainage from task-responsive visual regions, including fusiform cortex, rather than the amygdala itself [34]. Given that emotional face matching is among the most widely used paradigms to study amygdala response, these results raised major concerns that apparent amygdala activations in conventional fMRI studies might be confounded by vascular signal. Introduced by Hariri and colleagues, variants of the emotional face-matching task – including the Human Connectome Project Young Adult (HCP-YA) emotion task examined here – have been used in over 250 fMRI studies [35, 36]. Direct comparison and meta-analytic evidence further indicate that emotional faces elicit larger amygdala responses than other emotional visual stimuli [37, 38].

Since the original report a decade ago [34], three key questions have remained unresolved. First, there has been little attempt at replication or generalization of the original finding, leaving its critique of unclear validity. Second, it is in particular unclear whether the putatively venous signal reported is specific to the visual face- and scene-matching paradigms examined in [34], or whether the reported confound would generalize across other tasks and sensory modalities. Third, the source of the venous signal has yet to be fully identified; it may reflect task-evoked signal from fusiform cortex (as proposed in the original study [34]), from neighboring medial temporal structures, from the amygdala itself, or from some mixture of these territories.

In this study, we revisit the basal vein problem using large-sample Human Connectome Project task and resting-state fMRI data together with task fMRI and vascular precision imaging from an independent internal sample. We first ask whether the peri-amygdalar signal appears across a diverse set of tasks, including emotional-face processing, working memory, social cognition, and auditory story comprehension. We then investigate whether this signal is coincident with the basal vein and whether its timing and source profiles are more consistent with distal visual cortex alone or with contributions from the amygdala and other territories. We conclude that BOLD fMRI of the human amygdala is indeed confounded by venous signal, that the amygdala itself, when engaged, is the leading candidate contributor to this signal, and that other territories drained by the BVR contribute task-evoked signal as well. The peri-amygdalar signal is thus sensitive to but not specific for amygdala engagement.

## Results

We analyzed both task (*N* = 311) and resting-state (*N* = 305) fMRI datasets from the HCP-YA cohort together with an internal precision-imaging movie-watching dataset (Dense Amygdala, *N* = 3) [39], with additional task fMRI and vascular imaging performed in the same three participants. Participant characteristics, acquisitions, and paradigms are summarized in Supplementary Table S1.

### Amygdala-activating HCP-YA tasks reveal reproducible peri-amygdalar responses

Task conditions that evoked positive BOLD responses within the anatomical amygdala also revealed a bilateral response dorsomedial and posterior to the amygdala, approximately coincident with the crural and ambient cisterns and the vessels running within them, including the basal vein of Rosenthal. Mean canonical beta within the CIT168 amygdala mask was largest for language story (0.183), emotion fear (0.152), social mental (0.124), and working-memory 0-back faces (0.124) (Figure 1A). These tasks correspond to stimuli of auditory narratives, emotional faces, socially interacting shapes, and neutral faces, respectively. The corresponding control conditions were near zero or negative, including language math (-0.031), emotion shape (-0.008), social random (0.019), 0-back places (0.019), and 0-back tools (0.015). These task responses defined four primary contrasts for anatomic mapping: fear-shape, faces-places, mental-random, and story-math.

**Figure 1.**
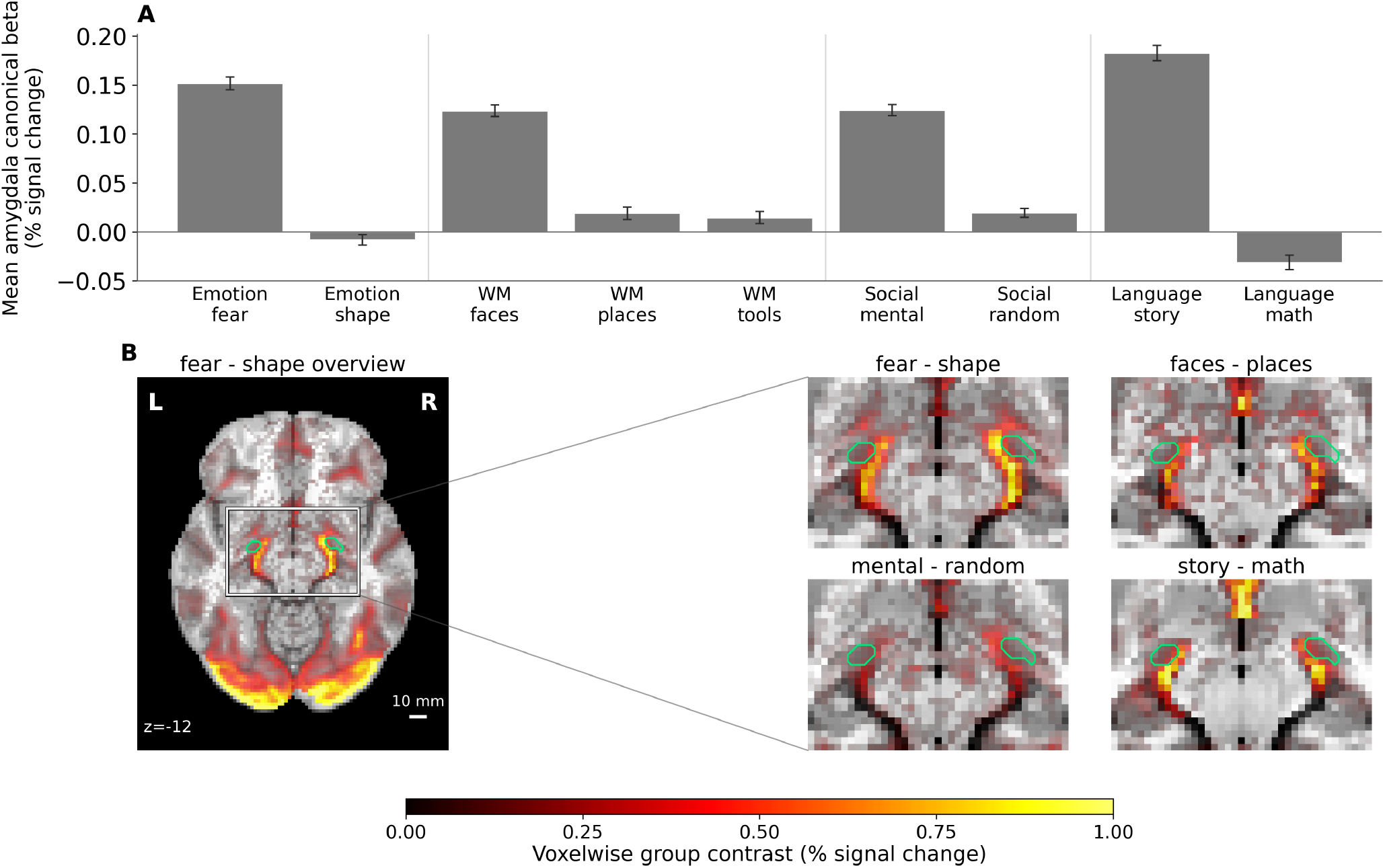
Amygdala-evoking HCP-YA tasks reveal a reproducible peri-amygdalar signal. **(A)** Mean amygdala canonical beta across task conditions, expressed in percent-signal-change units. Bars show group mean *±*SEM across participants (*N* = 311) of mean beta within the CIT168 anatomical amygdala mask. **(B)** Group task-contrast maps for emotion, working memory, social cognition, and language shown on the same axial slice (*z* = −12 mm). All positive group beta values are displayed on a shared color scale from 0 to 1.0% signal change, with opacity increasing with beta magnitude. The CIT168 anatomical amygdala mask is shown with a green outline. All images are unsmoothed with a native voxel resolution of 2 mm.

All four contrasts showed a similar medial peri-amygdalar pattern near the amygdala boundary, strongest for fear-shape and next strongest for story-math (Figure 1B). The story-math contrast is particularly informative because it produced the peri-amygdalar pattern without a visual stimulus. The recurring pattern therefore cannot be reduced to a face-specific display feature of the HCP-YA emotion task.

Group canonical-beta maps for all 23 modeled HCP-YA task conditions are provided in Supplementary Figure S1.

### The peri-amygdalar signal overlaps venous territory in group and subject-specific anatomy

The peri-amygdalar patterns derived from the four contrasts above overlapped the expected territory of the BVR in the vicinity of the amygdala. Anatomically, the BVR courses posteriorly from the union of the deep middle cerebral, anterior cerebral, and inferior striate veins through three primary segments to the vein of Galen, receiving tributaries from different territories along its course (Figure 2). The boundaries of these segments were originally proposed by Huang and Wolf [40, 41], which we here refer to as the striate (BVRs), peduncular (BVRp), and mesencephalic (BVRm). It should be noted that the BVR exhibits a large amount of individual variation in the presence and development of segments, including hypoplasia or aplasia, and in the specific drainage pathways to the galenic system and venous sinuses [42, 43].

**Figure 2.**
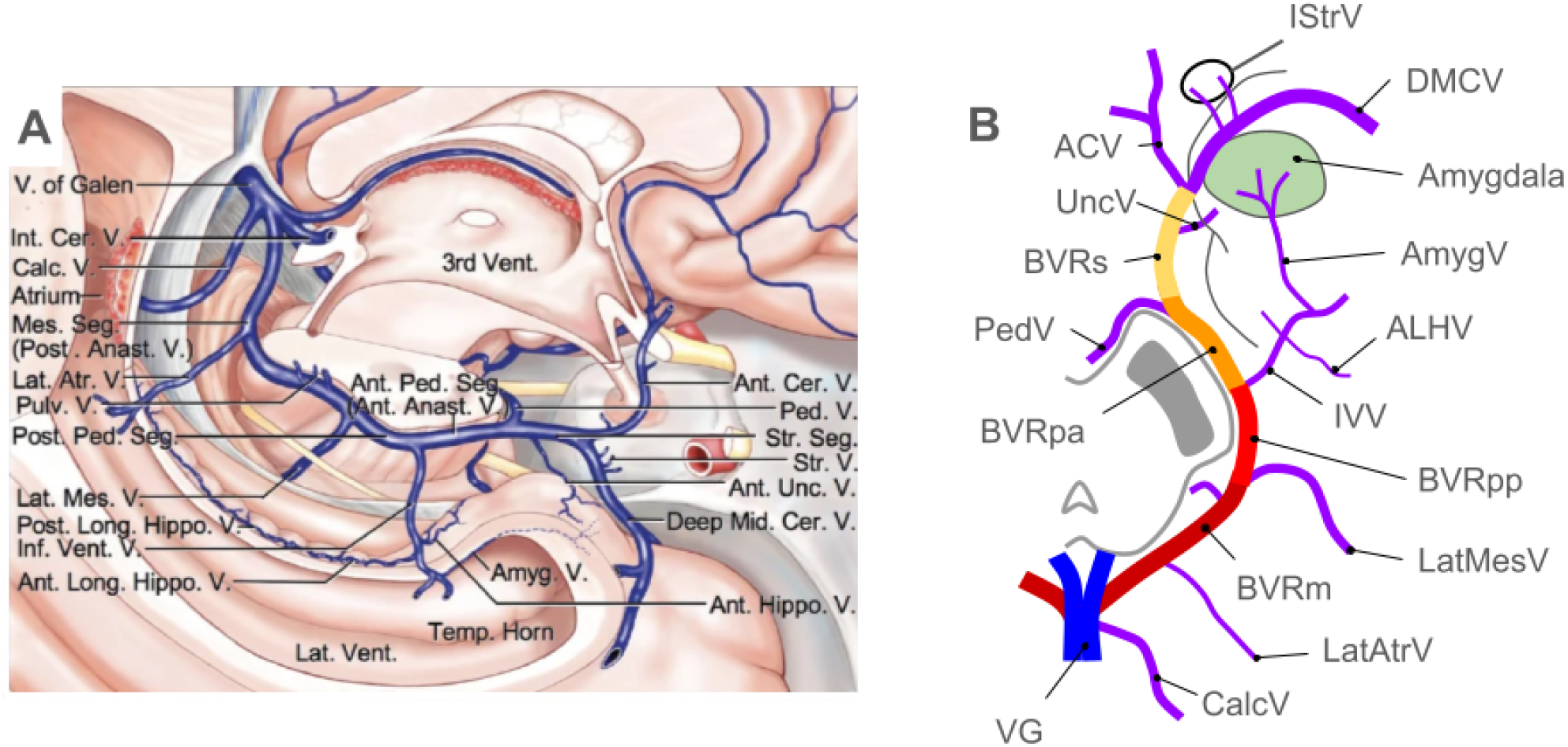
Anatomy and tributaries of the basal vein of Rosenthal. **(A)** Illustration of the typical course and anatomy of the basal vein, which most commonly drains anterior to posterior from its origin at the union of the deep middle cerebral vein (Deep Med Cer V), inferior striate vein (StrV) and anterior cerebral vein (Ant Cer V) to its final union with the vein of Galen (V of Galen). Note that the most posterior BVR segment is labeled here as mesencephalic, using the embryological convention. Many congenital variants of the BVR occur; this illustration represents the fully connected, typical variant of the BVR. Reproduced with permission from [26]. **(B)** Simplified schematic of basal vein showing primary tributaries and anatomical segments. The peduncular segment (BVRp) can be further divided into anterior (BVRpa) and posterior (BVRpp) segments at its union with the inferior ventricular vein (IVV). Venous outflow from the amygdala reaches the BVRp through the amygdalar vein (AmygV) and IVV, with possible auxiliary drainage to BVRs through the uncal vein (UncV) [25, 26]. Other abbreviations: IStrV - inferior striate vein, DMCV - deep middle cerebral vein, ACV - anterior cerebral vein, PedV - peduncular vein, ALHV - anterior longitudinal hippocampal vein, CalcV - calcarine vein, LatMesV - lateral mesencephalic vein, LatAtrV - lateral atrial vein, VG - vein of Galen.

In the HCP-YA data, where individual angiography was unavailable, the peri-amygdalar signal followed regions of high venous density in the population-level Venous Neuroanatomy (VENAT) atlas medial to the amygdala, while lying close to but largely outside the anatomical amygdala contour (Figure 3) [44].

**Figure 3.**
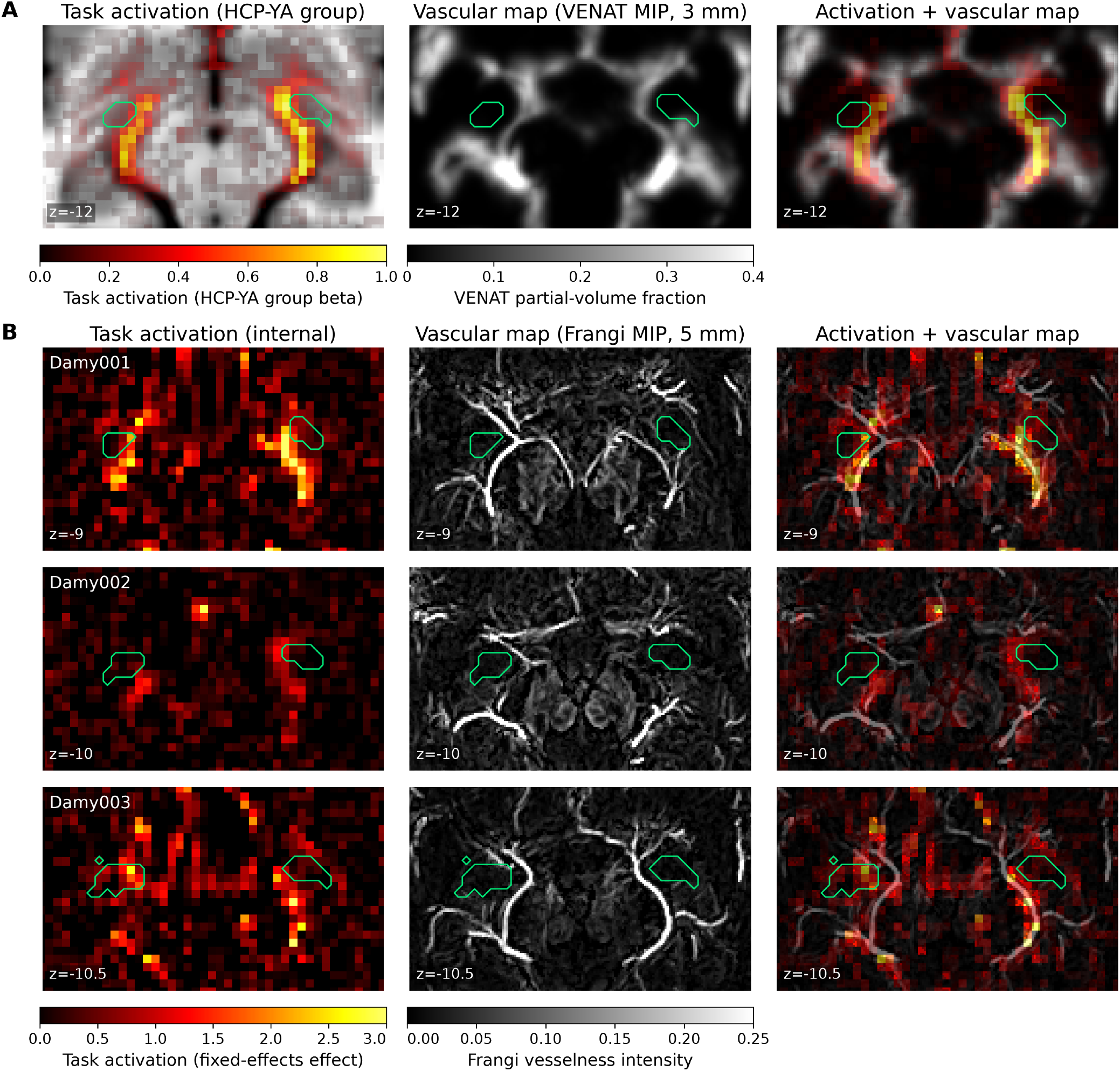
Task-evoked peri-amygdalar activation overlaps vascular anatomy. **(A)** HCP-YA group emotion-task activation for the fear-shape contrast (left), VENAT venous density (middle), and combined map (right). **(B)** Individual Dense Amygdala fear-shape fixed-effects activation maps (single 2-mm axial slices; left column), individual Frangi-filtered QSM venograms, each shown as a 5-mm axial maximum-intensity projection (MIP) centered at the corresponding level (middle), and combined maps (right). Coordinates are reported in MNI space in **(A)** and subject-specific space in **(B)**. Individual CIT168 amygdala boundaries are shown in green throughout.

The same pattern was visible at the individual-subject level in the Dense Amygdala dataset: in each participant, fixed-effects task activation formed a peri-amygdalar pattern that aligned with the BVR in subject-specific vascular maps (Figure 3). Fixed-effects maps of responses to the presence of faces, estimated from 6.20 hours of naturalistic movie viewing per participant, reproduced this alignment (Supplementary Figure S2). Expanded multimodal views combining subject-specific T2-weighted anatomy, time-of-flight (TOF) arteriography, quantitative susceptibility mapping (QSM)-derived venography, and CIT168 amygdala contours are provided in Supplementary Figure S3. Both venous and functional imaging results varied across the three participants with Damy002 showing the weakest task response overall, with BVRs and BVRp not identified bilaterally in the venogram.

### Amygdala task responses are organized by proximity to the striate BVR

Task-locked signal was present both inside the anatomical amygdala and in the striate and peduncular BVR ROIs. Averaged across the four activated HCP-YA task conditions, the anatomical amygdala peri-stimulus time course peaked at approximately 0.13% signal change, whereas the striate and peduncular BVR ROIs peaked at approximately 0.37% and 0.27% signal change, respectively (Figure 4B). Thus, while task-locked signal was present inside the amygdala proper, the peri-amygdalar BVR response was larger.

**Figure 4.**
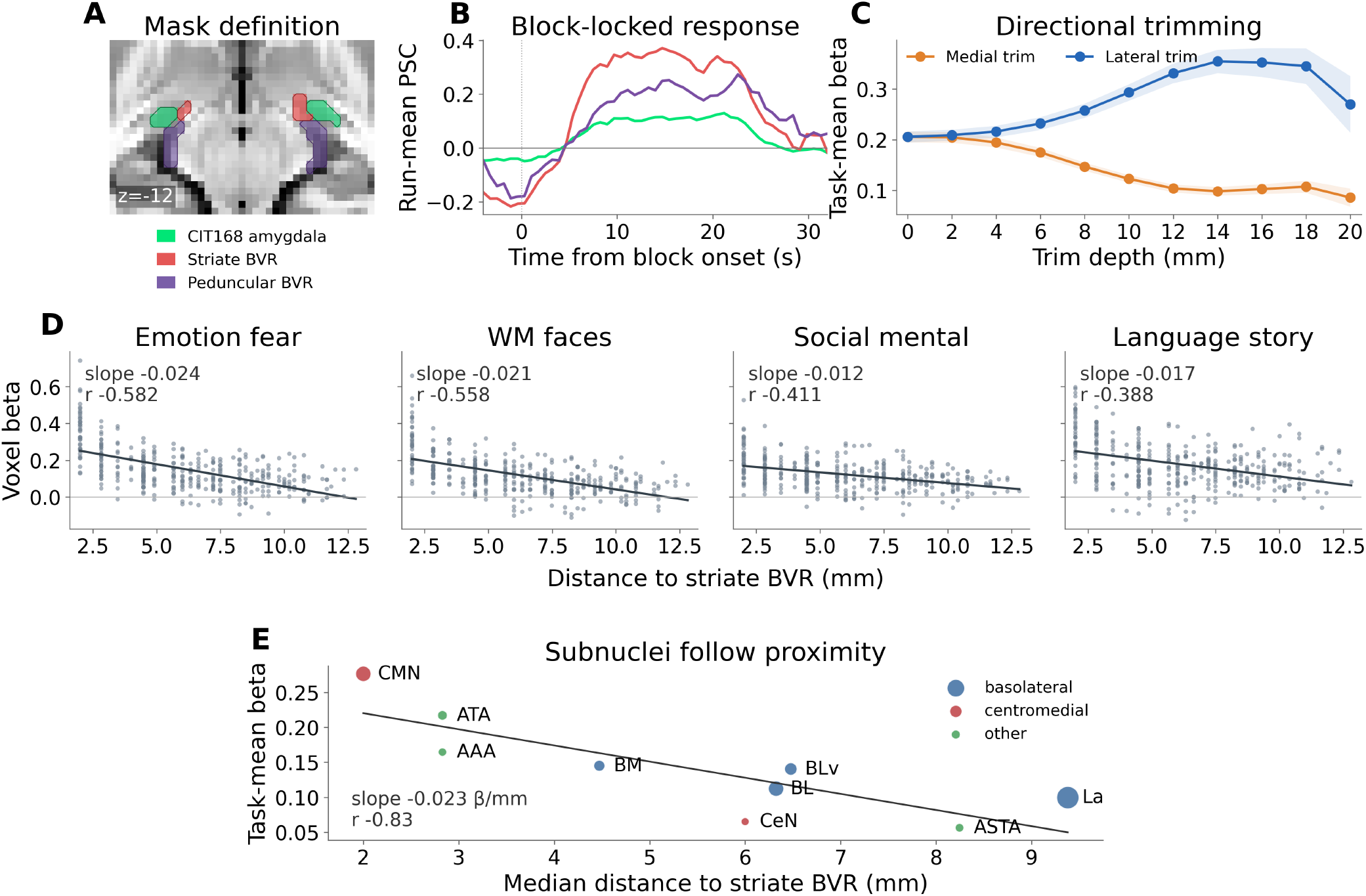
Amygdala task responses follow proximity to the striate BVR. **(A)** Anatomical amygdala and functionally defined BVR masks shown on the representative axial slice used for the spatial analyses. Green denotes the CIT168 anatomical amygdala, red denotes the adjacent striate BVR mask, and purple denotes the posterior peduncular BVR mask. **(B)** Block-locked percent-signal-change responses averaged across the four task conditions shown in Figure 1: emotion fear, working-memory faces, social mental, and language story. Responses are shown for the anatomical amygdala and the striate and peduncular BVR ROIs. **(C)** Directional trimming analysis of task-mean beta. Task-mean beta decreases rapidly with medial trimming and increases with lateral trimming, consistent with signal concentration in the medial, BVR-associated portion of the mask. **(D)** Voxelwise anatomical-amygdala beta as a function of distance to the adjacent striate BVR mask for each task condition. Points denote amygdala voxels; lines show ordinary least-squares fits. **(E)** Mean task beta over the four tasks for each CIT168 amygdala subnucleus plotted against its median distance from the striate BVR mask. Point size reflects the number of voxels in each subnucleus. This regression is descriptive; participant-level voxelwise distance slopes were the unit of inference.

Across the combined anatomical-amygdala and adjacent striate-BVR mask, task-mean beta was organized along the medial–lateral axis. Retaining progressively more medial voxels by trimming from the lateral side increased mean beta from 0.206 at zero trim depth to 0.355 at 14 mm. Removing the medial voxels closest to the striate BVR mask had the opposite effect, reducing mean beta to 0.086 at the deepest trim level (Figure 4C). This directional asymmetry indicates that signal across the combined mask is concentrated in its medial, BVR-associated portion.

The spatial gradient was also visible voxelwise. Group-median beta declined with distance from the striate BVR mask in all four task conditions, with slopes of -0.024 beta/mm for emotion fear, -0.021 beta/mm for working-memory faces, -0.017 beta/mm for language story and -0.012 beta/mm for social mental (Figure 4D). The same effect of distance is visible at the subnuclear level. The CIT168 corticomedial nucleus was the closest subnuclear group to the striate BVR mask and had the largest task-mean beta (Figure 4E). Subnuclear activation closely followed distance to the striate BVR mask and did not map neatly onto conventional functional groupings: the corticomedial (CMN, 2 mm from the mask) and central (CeN, 6 mm) nuclei of the same centromedial grouping fell at opposite ends of the response range in accordance with their distances.

### Spatial smoothing merges peri-amygdalar signal with amygdala parenchyma

Spatial smoothing transformed a narrow peri-amygdalar pattern into a broader amygdala-region effect. Internal participant Damy001 fear-shape fixed-effects z map is shown as an exemplar. While the unsmoothed map preserved spatially distinct peri-amygdalar signal adjacent to the amygdala contour (Figure 5A), increasing the smoothing kernel from 0 to 2, 4 and 6 mm FWHM progressively broadened this signal across the amygdala. At 4 and 6 mm FWHM – smoothing choices that are frequently adopted in the literature – the map no longer preserved a clear separation between the venous-territory signal and amygdala parenchyma. Conventional smoothing can therefore convert a spatially distinct peri-amygdalar pattern into an apparently ordinary amygdala activation map.

**Figure 5.**
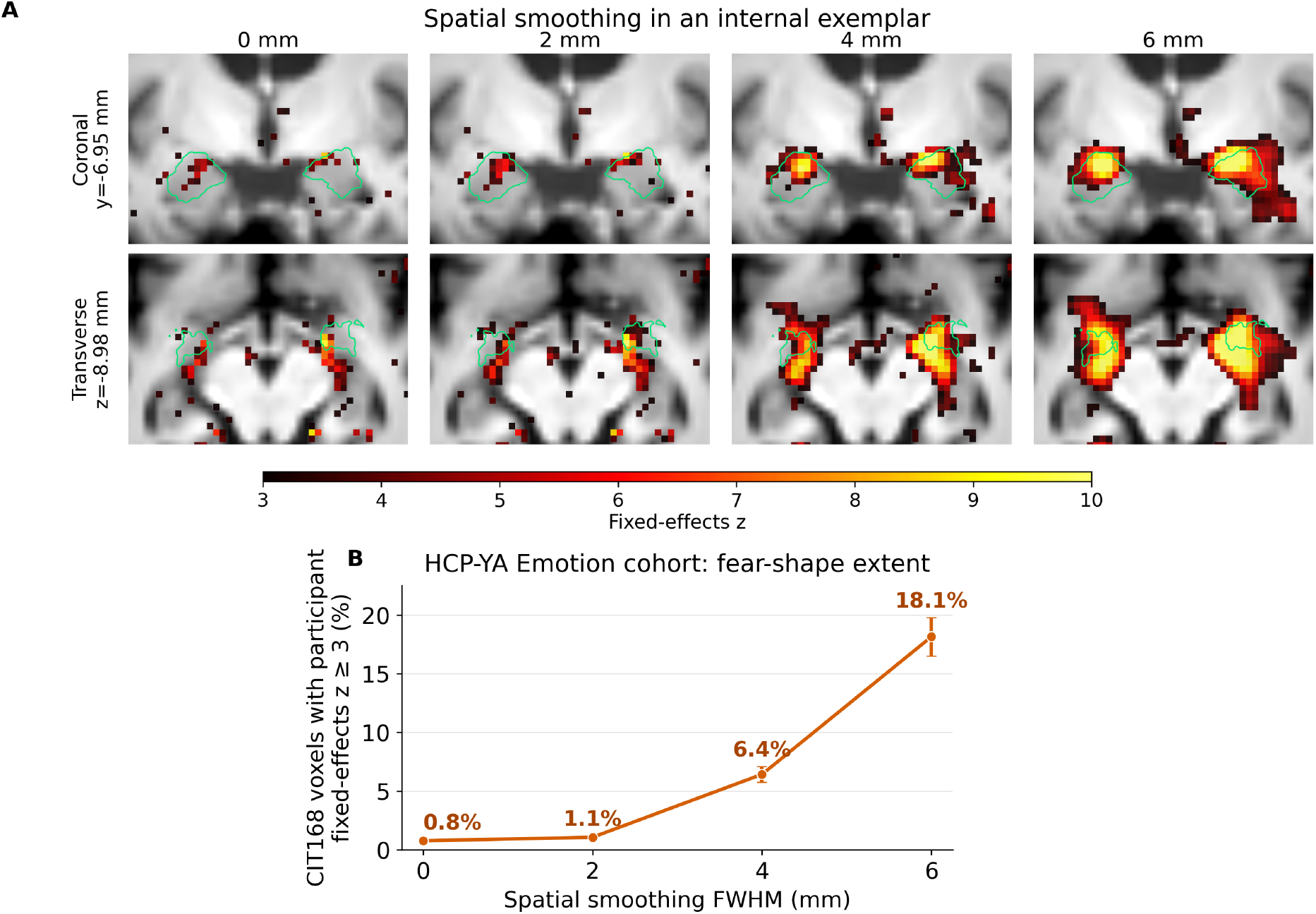
Spatial smoothing obscures the distinction between peri-amygdalar signal and amygdala parenchyma. **(A)** Coronal (*y* = −6.95 mm) and transverse (*z* = −8.98 mm) views of the Damy001 internal fear-shape fixed-effects z map after first-level GLM fitting with 0, 2, 4, or 6 mm spatial smoothing. Maps share a positive display range of *z* = 3 to 10. Across both views, increasing spatial smoothing makes the peri-amygdalar signal appear progressively less separable from the anatomical amygdala. The green contour denotes the subject-specific CIT168 amygdala mask derived from the probabilistic segmentation and thresholded at 0.5. **(B)** HCP-YA Emotion fear-shape results across the same smoothing levels. Points show the mean percentage of CIT168 amygdala voxels with participant fixed-effects *z* ≥ 3, and error bars show t-based 95% confidence intervals across participants (*N* = 311). The CIT168 mask was held fixed across smoothing levels. Mean amygdala beta remained between 0.160 and 0.168.

This effect generalized across the HCP-YA Emotion sample. The mean percentage of CIT168 amygdala voxels with fixed-effects *z* ≥ 3 increased from 0.8% without smoothing to 18.1% at 6 mm FWHM, while mean amygdala beta remained between 0.160 and 0.168 (Figure 5B). Smoothing therefore substantially expanded the apparent amygdala extent in z-statistic maps without generating a larger mean amygdala response.

### Resting-state lag structure supports amygdala-to-BVR temporal ordering

Resting-state timing separated signal in the BVR ROIs from the anatomical amygdala. To estimate relative timing, we used Rapidtide, which estimates voxelwise delays in the systemic component of low-frequency BOLD fluctuations. Because this component propagates with the circulation, it can provide a marker of relative vascular timing [45, 46, 47]. The striate BVR segment lagged the amygdala by a median of 1026 ms, and the peduncular BVR segment lagged the amygdala by a median of 1729 ms (Figure 6B). The striate segment was later than the amygdala in 94.4% of participants, and the peduncular segment was later than the amygdala in 97.0%. The peduncular segment was later than the striate segment in 84.9% of participants.

**Figure 6.**
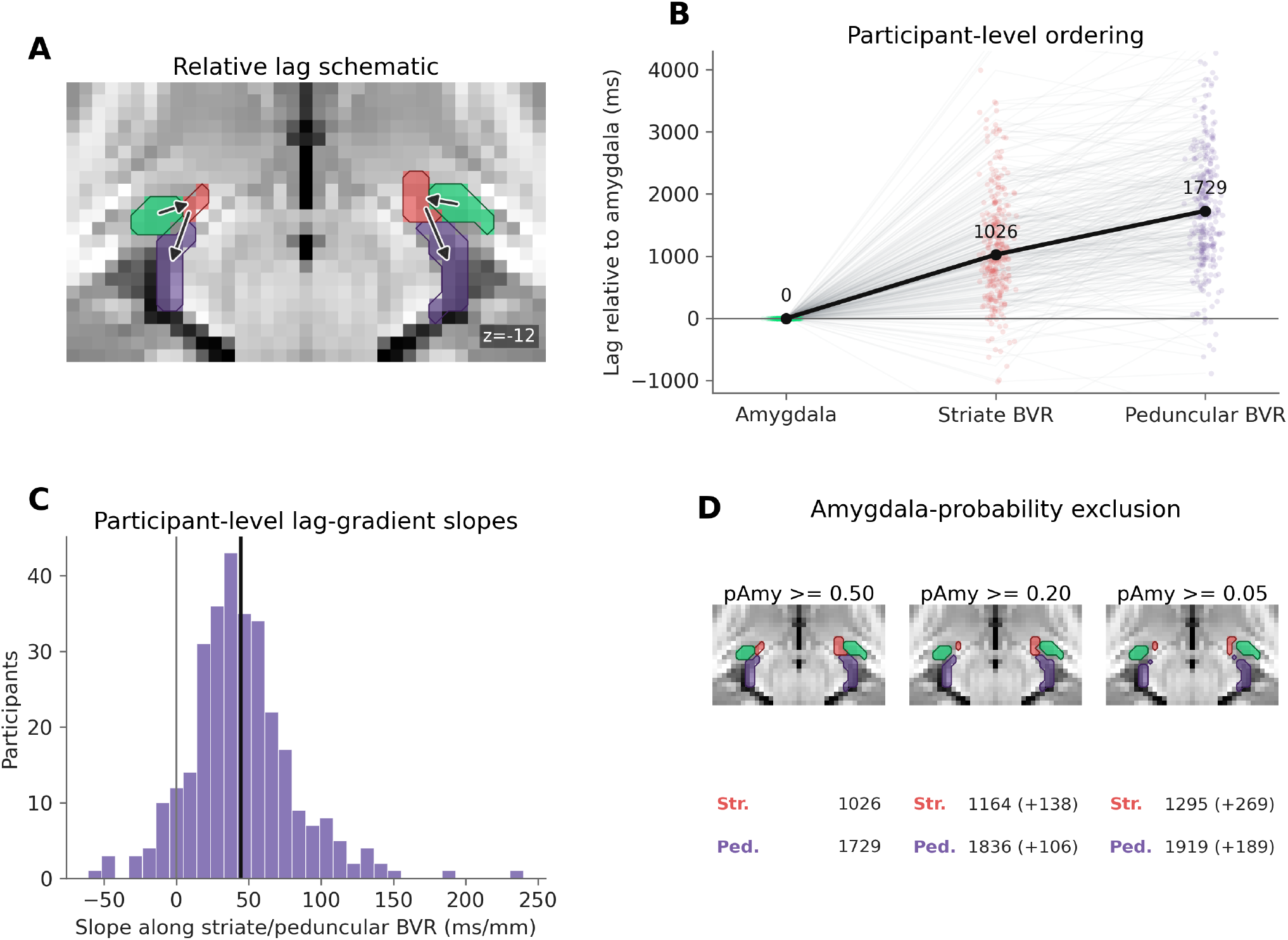
Resting-state lag mapping supports temporal separation between amygdala and the peri-amygdalar BVR signal. Rapidtide lag estimates were computed from minimally preprocessed resting-state fMRI, with participant-level consensus estimates across rest runs. **(A)** Anatomical schematic of the CIT168 amygdala mask (green), striate BVR segment (red), and peduncular BVR segment (purple) on the displayed axial slice. Arrows summarize the observed temporal ordering from amygdala to striate BVR to peduncular BVR. **(B)** Participant-level lag estimates relative to the amygdala. The black line shows the median over participants. **(C)** Distribution of participant-level lag-gradient slopes along the combined striate/peduncular BVR trajectory. The black vertical line shows the mean over participants. **(D)** Sensitivity to the anatomical amygdala boundary. BVR masks were recomputed after excluding voxels with CIT168 amygdala probability *p*_Amy_ ≥ 0.50, *p*_Amy_ ≥ 0.20, or *p*_Amy_ ≥ 0.05. Values show median lag relative to the amygdala.

Delay also increased along the combined striate/peduncular BVR trajectory. Participant-level lag-gradient slopes were positive in 91.5% of participants, with a mean slope of 44.5 ms/mm (Figure 6C). The combined gradient primarily reflects the striate-to-peduncular step together with a positive gradient within the longer peduncular segment (+126 ms/mm, positive in 90.5% of participants); the short striate segment showed no consistent internal gradient.

Stricter amygdala-overlap exclusions strengthened rather than weakened the lag effect. After removing voxels with *p*_Amy_ ≥0.20, median lags increased to 1164 ms for the striate BVR segment and 1836 ms for the peduncular BVR segment. After removing voxels with *p*_Amy_ ≥0.05, median lags increased to 1295 ms and 1919 ms, respectively (Figure 6D). These timing results place the BVR-associated signal downstream of the amygdala in a drainage-compatible sequence. The distinct timing profiles of the anatomical amygdala and two successive BVR segments demonstrate temporal separation of their signals, arguing against the peri-amygdalar signal being merely residual amygdala parenchyma outside an overly restrictive mask.

### Task profiles identify amygdala as the leading candidate contributor to the peri-amygdalar signal

While the preceding analyses localize the peri-amygdalar signal to venous territory, they do not identify which neural territories contribute to the task-evoked signal measured there. We therefore compared the peri-amygdalar task profile across 23 conditions from seven HCP-YA tasks with the corresponding profiles of 195 cortical and subcortical regions. Figure 7 highlights seven candidates chosen for their anatomical or functional relevance: anatomical amygdala, piriform cortex, orbitofrontal area 47m, rhinal cortex, anterior hippocampus, posterior insula, and fusiform face complex (FFC). FFC tests the distal visual-source hypothesis proposed in the original report [34]. The other candidates represent the amygdala and nearby medial-temporal regions, along with frontobasal and insular regions with known venous routes to the anterior BVR [25, 26, 48, 49].

**Figure 7.**
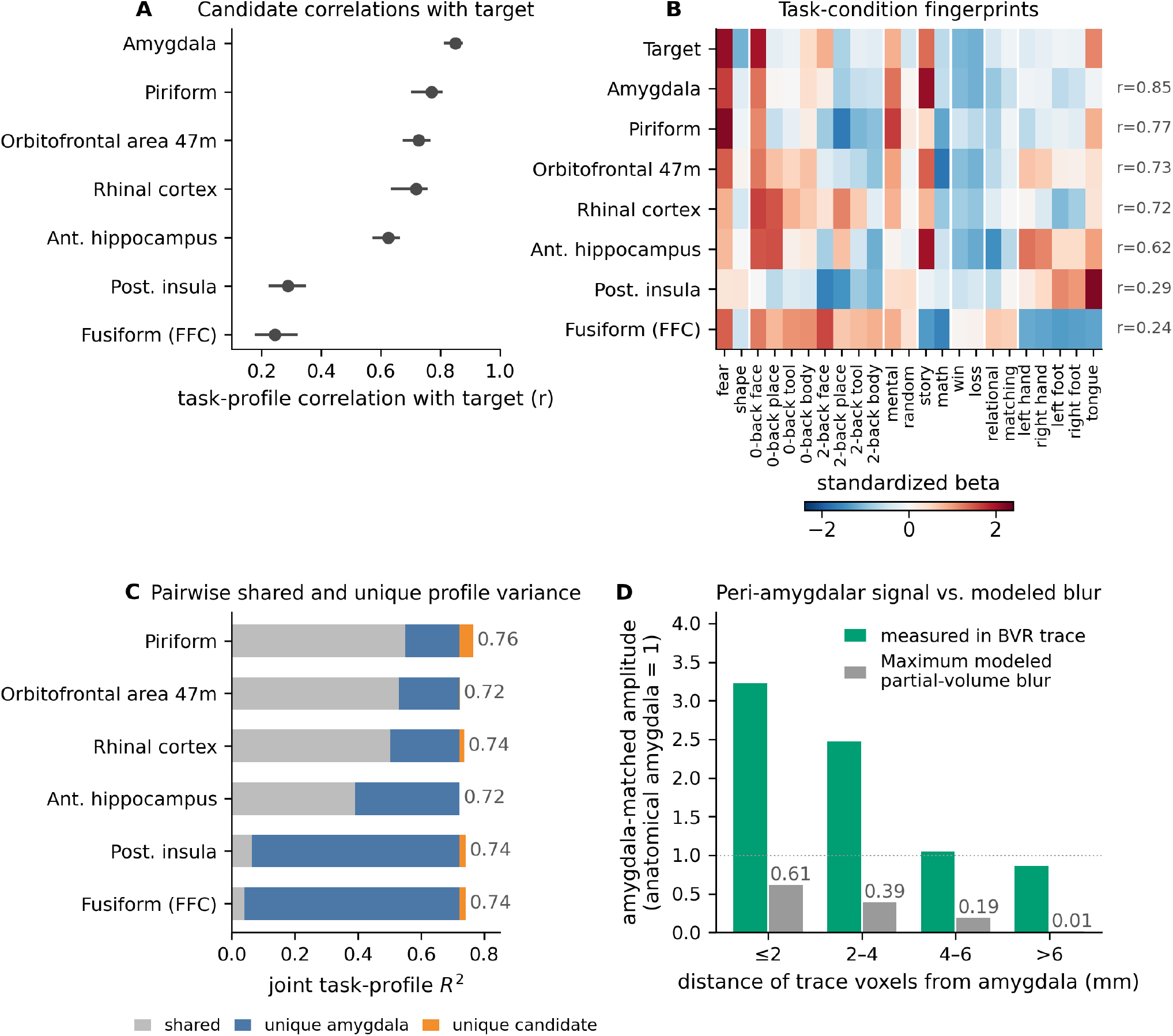
Task profiles support the amygdala as the leading candidate contributor to the primary peri-amygdalar target. All task-profile statistics give equal total weight to each of the seven HCP-YA task domains; whiskers show participant-bootstrap 95% confidence intervals (*N* = 311). **(A)** Group task-profile correlations between the primary peri-amygdalar target and the seven displayed candidate regions. The primary target combines the BVRs and BVRp segments after excluding voxels with CIT168 amygdala probability *p*_Amy_ ≥0.50; amygdala was the closest match among all 195 atlas regions. **(B)** Group mean fingerprints across the 23 task conditions for the target and the same candidates. Values are within-region standardized beta profiles (task-equal weights; hemispheres averaged for display); right-side labels give each candidate’s correlation with the target. Multi-parcel candidates average their standardized constituent profiles: the rhinal-cortex composite comprises entorhinal cortex (EC) and perirhinal ectorhinal cortex (PeEc), and posterior insula comprises the insular granular complex (Ig), para-insular area (PI), and posterior insular areas 1 and 2 (PoI1 and PoI2). **(C)** Pairwise variance decomposition for the amygdala and each of the other six candidates. Each bar partitions the joint *R*^2^ into shared variance (gray), variance unique to the amygdala (blue), and variance unique to the candidate (orange). These components describe model variance, not the proportion of venous signal arising from either territory. **(D)** Amygdala-matched signal amplitude in target voxels by distance from the amygdala (green), compared with the maximum modeled partial-volume blur of amygdala activity (gray; Gaussian kernels of 2–8 mm FWHM). Both are expressed relative to the anatomical-amygdala response, set to 1. The measured amplitude exceeds the blur prediction approximately 5.3-fold in the nearest shell.

For the primary peri-amygdalar target, which excluded voxels with CIT168 amygdala probability *p*_Amy_ ≥ 0.50, amygdala was the closest match among all 195 regions (*r* = 0.850, participant-bootstrap 95% CI [0.817, 0.870]; Figure 7A). Among the other displayed candidates, correlations were highest for piriform cortex (*r* = 0.769), orbitofrontal area 47m (*r* = 0.727), rhinal cortex (*r* = 0.719), and anterior hippocampus (*r* = 0.625); posterior insula (*r* = 0.288) and FFC (*r* = 0.244) were much weaker. The amygdala remained the top-ranked region when we omitted the two Emotion conditions used to define the target.

Stricter exclusion of voxels with probabilistic amygdala overlap weakened the amygdala match but did not materially alter the overall pattern. In the striate segment, the amygdala ranked first at the primary threshold and second to piriform at the two stricter exclusion thresholds (*p*_Amy_ ≥0.20 and ≥0.05); in the peduncular segment, it ranked first at all three thresholds (Supplementary Figure S5).

Pairwise models measured how much each of the six other displayed candidates added after accounting for the amygdala, and how much the amygdala added after accounting for that candidate (Figure 7C). For the primary target, the amygdala profile alone explained 72.2% of the variance in the group task profile. Adding an individual candidate to the amygdala-only model increased profile *R*^2^ by only 0.02–4.3 percentage points. In the reverse comparison, adding the amygdala to the corresponding candidate-only model increased *R*^2^ by 17.3–68.1 percentage points. Because the amygdala and piriform profiles were highly correlated (*r* = 0.77), most of the variance explained by piriform was shared with the amygdala. For each pair, these increments show how much one profile adds after accounting for the other; they do not estimate how much of the signal originated in either territory. As positive controls, the same procedure recovered the expected V1 profile for a V2 target and A1 profile for an LBelt–MBelt target (Supplementary Figure S4A).

The amygdala also provided the best single-region prediction of primary-target responses in held-out tasks (*Q*^2^ = 0.635). It remained the leading predictor when Emotion was excluded from both model fitting and evaluation (*Q*^2^ = 0.617).

Because the peri-amygdalar target borders the anatomical amygdala, partial-volume effects are a natural concern. In the point-spread models tested here, however, partial-volume contamination from the amygdala was too small to account for the amygdala-matched response in the primary peri-amygdalar target. Spatial blur can only dilute its source, yet the amygdala-matched response in voxels bordering the amygdala was 3.2 times the anatomical-amygdala response and exceeded the largest modeled blur prediction more than fivefold in every distance shell (Figure 7D; Methods). These results support a venous contribution beyond that predicted by simple partial-volume mixing, consistent with venous summation.

Using working-memory load, we next directly tested the hypothesis that the peri-amygdalar signal reflects drainage from distal visual areas such as the FFC. Consistent with prior HCP-YA findings [50], mean beta decreased from 0-back to 2-back faces in both the anatomical amygdala (0.124 to 0.037, *p* = 1.5 *×* 10^−24^) and the peri-amygdalar target (0.516 to 0.264, *p* = 1.4 *×* 10^−16^). By contrast, FFC increased (1.033 to 1.152, *p* = 9.4 *×* 10^−19^), while V1 was unchanged (0.470 to 0.474). At the participant level, the amount of suppression in the amygdala and target was also correlated (*r* = 0.41, *p* = 3.3 *×* 10^−14^).

Separately, a more posterior hotspot along the expected course of the BVRm had a different atlas-wide profile. It matched the posterior thalamus most closely, followed by a distributed set of cortical regions, including posterior visual, visuospatial, and inferior-temporal areas, and matched the amygdala only weakly (Supplementary Figure S4B,C). This pattern is consistent with different upstream contributions entering the BVR at different points along its course.

### The peri-amygdalar signal is sensitive to, but not specific for, anatomical-amygdala engagement

The source-profile analyses identified the amygdala as the leading candidate contributor to the primary peri-amygdalar target. We next asked whether the target response was specific to anatomical-amygdala engagement. Across all 23 HCP-YA task conditions, group mean canonical beta in the primary target tracked that in the CIT168 anatomical amygdala (*r* = 0.78; Figure 8A). This relationship remained similar after excluding the two Emotion conditions used to define the target (*r* = 0.76). Every condition with a clear positive anatomical-amygdala response showed a corresponding peri-amygdalar response.

**Figure 8.**
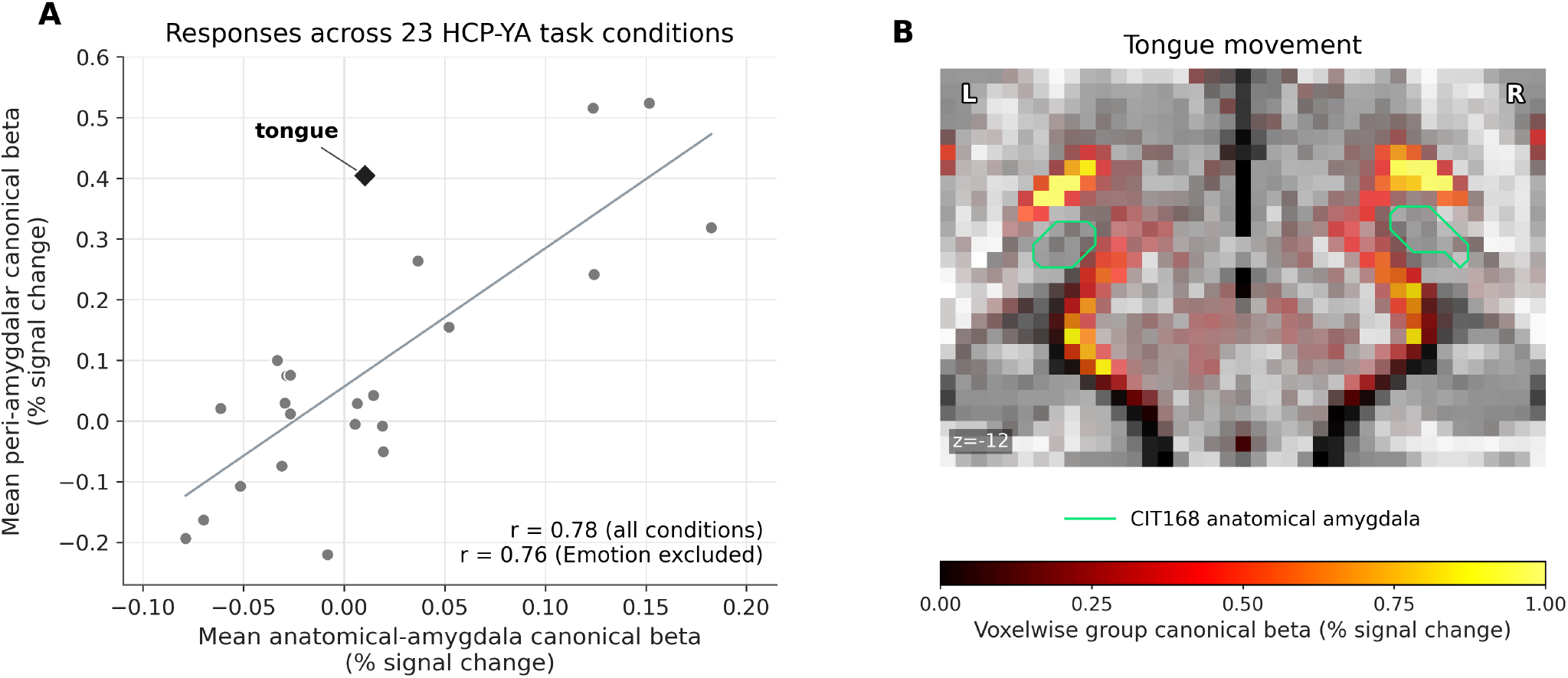
Across tasks, the peri-amygdalar signal is sensitive to, but not specific for, anatomical-amygdala engagement. **(A)** Group mean canonical beta in the CIT168 anatomical amygdala and primary peri-amygdalar target across the 23 HCP-YA task conditions (*N* = 311). Each point denotes one condition, and the line shows a descriptive ordinary least-squares fit. Responses were correlated across all conditions (*r* = 0.78) and after excluding the two Emotion conditions used to define the target (*r* = 0.76). Tongue movement (diamond) produced a prominent peri-amygdalar response despite a near-zero anatomical-amygdala response. **(B)** Unsmoothed group mean canonical-beta map for tongue movement relative to the implicit fixation baseline, shown at *z* = −12 mm. The green contour denotes the CIT168 anatomical amygdala.

Most conditions with little or no anatomical-amygdala response also showed little or no peri-amygdalar signal, with tongue movement as a clear exception. It produced a prominent response in the primary peri-amygdalar target (mean beta = 0.404% signal change) despite a negligible response within the anatomical amygdala (0.010%; Figure 8B). The tongue condition map also showed a separate bilateral hotspot anterior to the amygdala, near the ventral putaminal border and largely distinct from the primary peri-amygdalar target. Tongue movement recruits an extended orofacial network including the insula, frontal operculum, and putamen [51], and inferior striate veins drain parts of this network into the anterior BVR [48, 52]. This exception is consistent with the BVR acting as a shared venous collector: amygdala activation contributes signal when engaged, but task-evoked signal from other drained territories can enter the same pathway. Peri-amygdalar signal was therefore sensitive to, but not specific for, amygdala engagement.

## Discussion

In this study, we found that task-evoked fMRI near the amygdala contains a reproducible venous-territory component that can appear as focal parenchymal activation. The amygdala proper showed task-locked responses, but every condition that clearly engaged the anatomical amygdala also produced a corresponding larger peri-amygdalar signal. This signal aligned with both group-level and subject-specific vascular anatomy and followed a lag sequence at rest consistent with venous drainage. It persisted after excluding voxels with probabilistic amygdala overlap and became less spatially distinct from the anatomical amygdala after smoothing. The amygdala provided the closest task-profile match for the primary peri-amygdalar target, with rival candidates adding little profile information beyond it. Tongue movement provided an important boundary condition: it produced a prominent peri-amygdalar response despite negligible signal within the unsmoothed anatomical amygdala, showing that the peri-amygdalar signal is sensitive to, but not specific for, amygdala engagement. Together, these findings confirm Boubela et al.’s localization of peri-amygdalar signal to BVR territory but revise their favored interpretation that it was largely unrelated to amygdala activity and probably originated in fusiform cortex [34]. Amygdala activation contributes substantially to the peri-amygdalar signal when the amygdala is engaged, while the BVR also carries task-evoked contributions from other drained territories. Thus, broad claims about task engagement remain supported when based on unsmoothed signal within the CIT168 anatomical amygdala mask, but focal claims about precise parenchymal and subnuclear activation warrant considerable caution.

### Peri-amygdalar signal generalizes across tasks

The signal was not limited to visual or face stimuli: it appeared across several HCP-YA task domains and was among the strongest during auditory story comprehension, which evoked the peri-amygdalar pattern without visual input. It appeared even for tongue movement, plausibly via venous drainage from the broader orofacial network (Figure 8). Venous effects can preserve task information while displacing where it is measured [53]. The HCP-YA task analyses and the internal task and movie analyses all used ICA-FIX-cleaned data; the HCP-YA data additionally underwent spatial-ICA component reclassification and temporal-ICA cleanup. The persistence of the peri-amygdalar signal across these datasets indicates that modern ICA-based denoising does not remove this confound.

### These results change the spatial interpretation of amygdala fMRI

Amygdala parenchyma responded across emotional, social, and cognitive tasks, consistent with prior literature [3, 12, 38]. In the present data, however, apparent subnuclear differences followed proximity to the striate BVR mask, undermining attribution of these responses to precise locations or nuclei. A drainage-governed signal provides a plausible common mechanism for previously reported poor reliability, preprocessing sensitivity, and instability of subregional amygdala estimates [12, 13, 14, 15, 17, 54]: group-level engagement can remain reproducible even when the estimates required for subnuclear or individual-difference claims are not.

Spatial smoothing helps explain why this confound is easily overlooked: it merges a narrow BVR-associated signal with the amygdala in z-statistic maps, producing apparent activation across a much larger portion of the amygdala even though mean amygdala beta changes little (Figure 5). This confirms Boubela et al.’s concern that smoothing can convert a spatially distinct vascular pattern into apparent parenchymal activation [34].

### Venous-territory measurement and amygdala source are compatible

The peri-amygdalar signal’s venous location does not imply that the amygdala is uninvolved. At rest, the peri-amygdalar response followed the amygdala in a temporal sequence consistent with venous drainage. Across tasks, the amygdala was the closest match to the target profile among all 195 atlas regions, and the other displayed candidates added little target-profile variance once the amygdala profile was included. Modeled partial-volume contamination could not account for this result. Together, these findings support a vascularly displaced signal that reflects amygdala activity but may also include contributions from other drained territories.

This conclusion revises the simplest distant drainage account. Fusiform and visual regions can contribute task information to venous signals, and Boubela et al. highlighted that possibility for emotional-face tasks [34]. However, in the present source-attribution analysis – which successfully recovered upstream profiles in auditory and visual positive controls (Supplementary Figure S4) – the peri-amygdalar signal was not dominated by fusiform contributions. Further, working-memory load moved the target with the amygdala and opposite to FFC. Drainage anatomy also argues against a simple fusiform explanation. Basal temporal cortex generally drains through superficial or tentorial routes, whereas medial occipitotemporal outflow that reaches the BVR usually enters the posterior mesencephalic segment [25, 55]. The BVR most commonly drains anterior to posterior into the great vein of Galen (posterior outflow to Galen was observed on 87.8% of 500 sides in an angiographic series [43]), placing these posterior confluences downstream of the striate and peduncular segments when those segments are continuous. Even in the common variant lacking a striate–peduncular anastomosis, the striate segment drains anteriorly toward the cavernous or sphenoparietal sinus rather than receiving posterior inflow. Given individual variation in venous drainage we cannot exclude visual-cortex contributions, but neither the task profiles nor the anatomy favor them.

The typical anatomy of the BVR does, however, admit other contributors. Venous outflow from the amygdala reaches the peduncular BVR through the amygdalar and inferior ventricular veins, while superficial cortical amygdalar territory has an anatomically plausible route to the striate segment through anterior uncal veins [25, 26] (Figure 2B). The tributaries of the BVR at its anterior origin typically include the insular veins via the deep middle cerebral vein, inferior striate, anterior cerebral and olfactory veins [25, 26, 48, 49, 52]. Together, these tributaries provide plausible inflow to the striate BVR from anterior medial-temporal, insular, posterior basal-frontal, and selected inferior striatal territories. Piriform cortex matched the target closely, especially under strict amygdala exclusion, but this result requires caution: the HCP-MMP1 authors observed strong vascular artifacts in the piriform parcel and considered its results suspect [56]. The distinct posterior-thalamic and distributed cortical profile of the BVRm hotspot, with a substantial visual and visuospatial component, is likewise consistent with different upstream territories contributing at different points along the BVR (Figure 2B and Supplementary Figures S4 and S6). Taken together, the results support the amygdala as a principal but not exclusive contributor to the peri-amygdalar signal.

### Implications

Because broad claims of engagement remain supported while spatially specific claims do not, studies localizing effects to amygdala subnuclei or focal parenchymal clusters near the BVR should minimize smoothing and inspect subject-specific vascular anatomy when possible. A precise probabilistic anatomical amygdala atlas, such as CIT168, should also be used because broader atlas definitions can encompass more adjacent vascular territory (Supplementary Figure S7). When smoothing is used, researchers should inspect unthresholded effect maps and mean ROI effect estimates alongside z-statistic maps. Taken together, these safeguards improve interpretation but cannot resolve vascular mislocalization.

We emphasize that this is a claim about measurement, not about neurobiology. A functional differentiation among amygdala subnuclei is well established, but on evidence that does not share the hemodynamic basis of the present confound. Direct single-neuron recordings show face- and emotion-selective responses in the primate amygdala [57] and in the human amygdala [58], and decades of rodent and primate circuit work assign distinct roles to the basolateral input and central output nuclei in defensive learning [59, 60, 61]. Human lesion studies support the same functional distinction: selective basolateral damage alters context-dependent face processing [62]. Our results do not challenge the functional differences among amygdala subdivisions. They show instead that conventional BOLD fMRI cannot reliably assign signal to those subdivisions.

### Limitations and Future Directions

All fMRI analyzed here was acquired at a field strength of 3 Tesla, so direct generalization to ultra-high-field MRI protocols remains to be tested, although gradient-echo BOLD remains sensitive to draining-vein signal at higher field [27, 53]. VENAT is a population atlas and cannot capture individual venous variants. Likewise, the three internal participants demonstrate subject-specific anatomical alignment but cannot establish how common a given pattern is. Larger cohorts combining task fMRI with venography are therefore needed. Performing QSM venography at higher field strengths such as 7 Tesla allows for both higher spatial resolution and greater T2^*^ sensitivity [63, 64], allowing direct detection of smaller veins which may contribute to peri-amygdalar task-correlated signal. The use of T2-prepared GRE or complex-valued EPI may improve separation of large and small vessel BOLD signal, as has been demonstrated for high field layer specific fMRI [33, 65, 66]. Source attribution remains indirect. Task-profile similarity and incremental variance help narrow the plausible contributors, but they cannot establish where the signal arose or how much each territory contributed. More direct approaches, such as the recently developed perfusion source mapping, may help answer these questions [67].

## Methods

Structural and functional MRI data were analyzed from two sources: a retrospectively selected subsample of the Human Connectome Project Young Adult (HCP-YA) release and three prospectively studied participants from the internal Dense Amygdala cohort.

### Human Connectome Project Young Adult data

Derived data were drawn from the HCP-YA cohort [68]. Functional images were acquired on a customized 3 T Connectome Skyra (Siemens Medical Solutions, Malvern, PA) using multiband gradient-echo EPI at 2 mm isotropic resolution and TR = 720 ms; imaging parameters and task paradigms are described in detail elsewhere [5, 69, 70]. Each task comprised one left-right (LR) and one right-left (RL) phase-encoded run. Resting-state acquisitions comprised four 1,200-volume runs (REST1 and REST2, each acquired with LR and RL phase encoding), resulting in approximately 58 minutes of imaging data per participant. The analyzed acquisitions and conditions are summarized in Supplementary Table S1.

Participants from the HCP-YA cohort were included in this analysis if complete 3 T task fMRI was available, no HCP-YA quality-control issue was recorded, and task accuracy was at least 80% for the Emotion, Language, Relational, and Working Memory tasks. Of 316 participants meeting these criteria, five lacked complete task inputs, leaving a final task sample of *N* = 311. Resting-state analyses included the 305 participants with all four resting-state runs.

Task models used the 2025 HCP-YA Task3T Recommended volumetric derivatives. Core spatial preprocessing followed the HCP-YA minimal-preprocessing framework [71]; the 2025 derivatives additionally incorporate spin-echo-based bias-field correction, multi-run ICA-FIX, spatial-ICA component reclassification (Reclean), and temporal-ICA cleanup [72]. No additional nuisance regressors were included, and no spatial smoothing was applied in the primary task models. The S1200 release was used only for the resting-state lag analysis.

### General linear modeling of HCP-YA tasks

Emotion, Gambling, Language, Motor, Relational, Social, and Working Memory runs were modeled separately using Nilearn v0.13.1 [73]. Events were read from the HCP-YA three-column EV files. The Motor cue was included as a nuisance condition but excluded from scientific contrasts and task profiles. Each run was fitted with an SPM canonical hemodynamic response function and its temporal derivative, a cosine drift model with a 1/128-Hz high-pass cutoff, and an AR(1) noise model. Voxelwise signal was scaled to a run mean of 100, giving the canonical effects percent-signal-change units.

The LR and RL run estimates were combined within participant using inverse-variance-weighted fixed effects. Group display maps were calculated as the voxelwise mean of participant fixed effects. These maps are descriptive and are not formal second-level tests. ROI analyses used the participant as the unit of analysis.

### Amygdala and peri-amygdalar masks

We defined the anatomical amygdala from the CIT168 probabilistic amygdala atlas [74]. The bilateral probability map was resampled to the 2-mm HCP-YA MNI152NLin6Asym grid using sinc interpolation and thresholded at *p*_Amy_ ≥0.50.

We defined the peri-amygdalar mask from the full-cohort Emotion fear-shape group effect map. Voxels with group beta greater than 0.5 were retained within six bilateral bounding boxes, defined by inspection of the group beta images to correspond approximately to the striate (BVRs), peduncular (BVRp), and mesencephalic (BVRm) segments. The striate and peduncular components formed the primary target and were also analyzed separately; the BVRm component was used for the supplementary source-profile analysis. The primary striate and peduncular masks excluded voxels with *p*_Amy_ ≥0.50, and exclusions at *p*_Amy_ ≥0.20 and *p*_Amy_ ≥0.05 tested sensitivity to the anatomical amygdala boundary. We used the VENAT partial-volume vein atlas [44], resampled to MNI152NLin6Asym space, as an independent population-level reference.

### Task-locked and spatial-gradient analyses

We extracted mean BOLD time series from the anatomical amygdala and the striate and peduncular BVR masks, converted them to percent signal change relative to the run mean, and sampled them from 4 seconds before through 32 seconds after block onset. Blocks were averaged first within participants and then across participants. Figure 4 shows the response averaged across Emotion fear, Working Memory 0-back faces, Social mental, and Language story.

To characterize spatial organization within the amygdala, we used two complementary analyses. First, we progressively trimmed the combined anatomical-amygdala and striate-BVR mask from its medial or lateral edge in 2-mm increments and measured mean canonical beta at each depth. Second, the Euclidean distance from each anatomical-amygdala voxel to the nearest striate-BVR voxel was calculated after excluding mask overlap. For each participant and each of the four Figure 4 task conditions, canonical beta was regressed on distance; participant-level slopes were the unit of inference. CIT168 subnuclear summaries were descriptive overlays on this distance analysis.

### Spatial-smoothing sensitivity

To quantify whether the Figure 5 exemplar generalized to HCP-YA data, the two HCP-YA Emotion runs were fitted at 0-, 2-, 4-, and 6-mm Gaussian FWHM with all other model settings held fixed. Fear-shape fixed-effects beta, variance, and z statistic were calculated within the fixed CIT168 amygdala mask. Apparent amygdala extent was summarized for each participant as the percentage of mask voxels with fixed-effects *z* ≥3; group means and t-based 95% confidence intervals were calculated across participants. Mean ROI beta was summarized at each level to distinguish a change in apparent z-map extent from a change in mean effect magnitude.

### Resting-state lag analysis

Resting-state data were used for the timing analysis because the four HCP-YA rest runs provided approximately 58 minutes of acquisition per participant, compared with 4.5 minutes across the two HCP-YA emotion runs [5, 70]. This greater data quantity is particularly valuable for estimating delays in low-frequency BOLD fluctuations, for which sampling error decreases strongly as the amount of data increases [75].

Resting-state lag mapping used the minimally preprocessed S1200 MNI-space volumes rather than the ICA-FIX-cleaned derivatives [76]. Rapidtide 3.1.6 was run separately on each of the four rest runs using delay-mapping mode with its default spatial smoothing and delay refinement, the low-frequency-oscillation filter preset, a −10 to +10 second lag-search range, and 10,000 null correlations [45]. The first 72 volumes were omitted, and the six rigid-body motion parameters were included as nuisance regressors without motion derivatives.

Run-level delay estimates were retained where they passed Rapidtide’s correlation-fit validity mask. For each participant, the consensus delay map was the voxelwise median across valid run-level maps. Subsequent ROI summaries required at least two valid runs per voxel. Regional timing was summarized as the median consensus delay within each mask. Lag relative to amygdala was calculated by subtracting the anatomical-amygdala median from the corresponding BVR-segment median. The spatial lag gradient was estimated from median delay in six distance bins along the combined striate–peduncular BVR mask.

### Dense Amygdala dataset

Precision functional MRI was acquired in three adult volunteers during approximately 520 minutes of passive movie watching. All data were acquired at the Caltech Brain Imaging Center using a 3 T whole-body MRI system (Prisma.Fit, Siemens Medical Solutions, Malvern, PA) equipped with a 32-channel phased-array head coil. T2^*^-weighted functional images were acquired from an approximately 60 mm oblique axial slab using the following imaging parameters: TR = 556 ms, TE = 30 ms, flip angle = 48°, isotropic voxel size = 2.0 mm, no in-plane acceleration, multiband slice acceleration = 4, and complex-valued reconstruction. Full acquisition details are reported in [39]. In addition to the original precision acquisitions, two whole-brain 3D gradient-recalled echo (GRE) time-of-flight (TOF) arteriograms were acquired in each participant without contrast enhancement using the following parameters: TR = 21 ms, TE = 3.4 ms, flip angle = 18°, voxel size = 0.26 *×* 0.26 *×* 0.5 mm, and superior spatial saturation. The two TOF acquisitions were affine-registered to each participant’s 0.5-mm subject-specific T1-weighted template, brain-masked, and averaged for visualization. Complex-valued whole-brain 3D multi-echo GRE images were acquired for quantitative susceptibility mapping with the following parameters: TR = 35 ms, TE = 5.3–31.5 ms in 3.74 ms increments, eight unipolar echoes, flip angle = 15°, and voxel size = 0.8125 *×* 0.8125 *×* 0.75 mm. Quantitative susceptibility maps were calculated using the QSMxT v8.2.1 pipeline [77], with ROMEO phase unwrapping [78] and rapid two-step dipole inversion [79], then affine-registered to the same 0.5-mm subject-specific template using the multi-echo magnitude image. Venograms were derived from the registered QSMs with a positive-ridge multiscale Frangi filter at six logarithmically spaced scales spanning 0.5–2.5 mm on the display grid [80]. The resulting vesselness-filtered images were gamma-corrected with exponent 0.5 to retain the visibility of both small, low-intensity veins and large, high-intensity veins without clipping the latter. The follow-up emotion-task fMRI, TOF angiography, and multi-echo GRE acquisitions were approved by the Caltech Institutional Review Board (protocol IR24-1490), and all participants provided written informed consent.

The four internal Emotion runs were modeled using the same unsmoothed GLM framework as the HCP-YA task analyses, and fear-shape estimates were combined across runs within participant. For Supplementary Figure S2, a binary face-presence regressor derived from framewise annotations was fitted to 29 unsmoothed ICA-FIX-cleaned movie runs per participant, and the estimates were combined within participant.

### Task-profile source-attribution analysis

Task profiles were constructed from the fixed-effects canonical beta for each of 23 HCP-YA task conditions in each hemisphere, yielding 46 condition-by-hemisphere cells per region. For each participant, we defined each cell as the median beta across the region’s voxels in that hemisphere, providing a robust summary for small, heterogeneous regions. Results were similar with voxelwise means: the primary target’s correlation was 0.78 with the amygdala and 0.74 with piriform cortex, leaving their ranking unchanged. We obtained the group profiles by averaging each cell across the available participants (up to 311). Because the number of modeled conditions differed across tasks, profile correlations and regressions used task-equal weights: each of the seven task domains contributed one seventh of the total weight, with equal weight assigned to conditions and hemispheres within each task.

We compared the primary and stricter-sensitivity peri-amygdalar targets defined above with 195 atlas-defined regions: CIT168 amygdala [74], 180 HCP-MMP1 cortical areas represented bilaterally [56], and 14 non-amygdala regions from the Tian S2 atlas [81]. Parcels from each atlas were represented in MNI152NLin6Asym volumetric space, resampled to 2-mm resolution. HCP-MMP1 parcels were derived from the publicly available group registration-fusion probability maps [82]. Each candidate region was intersected with a gray-matter support mask. This support mask was defined as the union of voxels classified as predominantly gray matter by FSL FAST [83] in the 2-mm MNI152NLin6Asym template and voxels labeled as subcortical gray matter in the standard HCP segmentation distributed through TemplateFlow [84].

The supplementary BVRm target comprised the 33 voxels outside the Tian S2 posterior-dorsal thalamus parcel and was compared with the same 195 candidate definitions. Before profile extraction, voxels overlapping the target were removed from each candidate mask so that direct overlap could not inflate target–candidate correlations. Figure 7 shows anatomical amygdala; HCP-MMP1 piriform cortex; orbitofrontal area 47m; rhinal cortex (EC and PeEc); Tian S2 anterior hippocampus; posterior insula (Ig, PI, PoI1, and PoI2); and HCP-MMP1 fusiform face complex. These candidates test four hypotheses about upstream source territory: anterior medial-temporal, frontobasal, insular, and distal visual. For candidates comprising multiple parcels, we standardized each parcel’s 46-cell profile using task-equal weights, then averaged the standardized profiles across parcels. We measured target–candidate similarity with weighted Pearson correlation, deriving confidence intervals from 1,000 bootstrap draws of participants.

For Figure 7C and Supplementary Figure S4A, three models were fit for each candidate: amygdala only, candidate only, and both together. Each region’s unique variance was the drop in *R*^2^ when it was removed from the joint model, with the remainder of the joint *R*^2^ classified as shared variance. The Supplementary Figure S4A controls paired an HCP-MMP1 V2 target with V1 and a standardized composite of HCP-MMP1 LBelt and MBelt with A1; these comparisons assess recovery of expected upstream-system profiles rather than causal source direction.

Using the same group profiles and masks, we fitted task-weighted linear models with an intercept to six tasks and predicted the seventh, repeating across tasks. Prediction accuracy was summarized as *Q*^2^ = 1 − SSE*/*SST, using task-weighted sums of squared prediction errors (SSE) and squared deviations from the overall target mean (SST). The Emotion-excluded analysis used five training tasks and one test task.

For Figure 7D, we used linear regression to estimate how much the response in each voxel of the primary peri-amygdalar target varied with the hemisphere-matched amygdala response across tasks. We expressed this amplitude relative to the anatomical-amygdala response. To model partial-volume blur, we smoothed the amygdala probability map with 2–8 mm FWHM Gaussian kernels. We normalized each prediction to the modeled response in the same amygdala reference region and took the largest value at each target voxel before averaging by distance shell.

Because the Emotion task was used to define the peri-amygdalar mask, we repeated the atlas screen without its two conditions to test whether the amygdala’s ranking depended on them.

For Figure 8A, we calculated the unweighted Pearson correlation across the 23 conditions between group mean canonical beta in the bilateral CIT168 amygdala and the primary peri-amygdalar target. We repeated the correlation after excluding the two Emotion conditions used to define the target.

For Supplementary Figure S5, we tested sensitivity to the amygdala boundary at three amygdala-overlap exclusions (*p*_Amy_ ≥0.50, ≥0.20, and ≥0.05) for the combined peri-amygdalar target and its striate and peduncular segments. For each target, we calculated the amygdala-profile correlation, the amygdala’s atlas rank, and the proportion of bootstrap samples in which it ranked first.

Supplementary Figure S6 displays the group emotion fear-shape map with the Figure 1 conventions on the *z* = −12 mm axial plane, on right-hemisphere sagittal slices at *x* = +14, +18, and +22 mm, and as maximum-intensity projections over a right-hemisphere sagittal slab (*x* = +6 to +30 mm) and a ventral axial slab (*z* = −28 to +2 mm), each over the single-slice anatomical background at the slab center. Segment arrows were placed within the corresponding BVR segment masks.

We assessed working-memory load using each participant’s fixed-effects betas for the 0-back and 2-back face conditions, averaged within the CIT168 amygdala, the primary peri-amygdalar target, and bilateral FFC and V1 masks. The FFC and V1 masks came from HCP-MMP1 maximum-probability labels thresholded at 20%. Load suppression was defined as 0-back minus 2-back response, tested against zero in each region using one-sample t tests. Across participants, we calculated the Pearson correlation between amygdala and target suppression values.

### AI Usage

No generative AI was used in the original conception of this study or in data collection. GPT-5.6 Sol and Claude Fable 5 assisted with refining and reviewing code and assembling the public release GitHub. These models also assisted with critiquing manuscript drafts. Z.D. reviewed all AI-assisted code, and all authors reviewed the resulting manuscript changes.

## Data availability

HCP-YA task and resting-state data are available from the Human Connectome Project (db.humanconnectome.org) subject to the HCP data use terms. The Dense Amygdala dataset is available through its OpenNeuro repository as described in [39]; the follow-up emotion-task, TOF, and QSM acquisitions analyzed here are available upon reasonable request to the authors.

## Code availability

All code required to reproduce the analyses and figures is available at github.com/zdiamandis/amyvasc-release.

## Acknowledgements

Data were provided in part by the Human Connectome Project, WU-Minn Consortium (Principal Investigators: David Van Essen and Kamil Ugurbil; 1U54MH091657) funded by the 16 NIH Institutes and Centers that support the NIH Blueprint for Neuroscience Research, and by the McDonnell Center for Systems Neuroscience at Washington University. This work was funded in part by an award from the Caltech Chen Center for Social and Decision Neuroscience.

## Author contributions statement

Z.D. analyzed the neuroimaging data and drafted the manuscript. J.M.T. acquired the MRI and behavioral data, analyzed the neuroimaging data, and drafted the manuscript. R.A. drafted the manuscript. R.A. and J.M.T. supervised the project and revised the manuscript critically for important intellectual content. All authors approved the final version to be published.

## Additional information

The authors declare no competing interests.

