## Supplementary Information for "Large-Vessel Venous Signal Confounds Apparent Amygdala Activation in BOLD fMRI"

**Table S1. Participant characteristics and analyzed MRI data. (A)** Participant characteristics. **(B)** HCP-YA task and resting-state fMRI. **(C)** Internal structural, functional, and vascular MRI.

**A. Participant characteristics**

| Dataset | *N* | Female | Male | Age, years |
|---|---|---|---|---|
| HCP-YA task cohort | 311 | 160 | 151 | 22–25 (82); 26–30 (146); 31–35 (79); 36+ (4) |
| HCP-YA rest subset | 305 | 157 | 148 | 22–25 (81); 26–30 (142); 31–35 (78); 36+ (4) |
| Internal Dense Amygdala | 3 | 1 | 2 | 28, 58, 62 |

**B. HCP-YA imaging data**

| Paradigm | Included conditions | Runs | *N* |
|---|---|---|---|
| Emotion | Emotional faces, shapes | 2 (LR/RL) | 311 |
| Working memory | 0-back and 2-back: faces, places, tools, bodies | 2 (LR/RL) | 311 |
| Social cognition | Mental and random shape animations | 2 (LR/RL) | 311 |
| Language | Auditory story, math | 2 (LR/RL) | 311 |
| Gambling | Win, loss | 2 (LR/RL) | 311 |
| Relational | Relational, matching | 2 (LR/RL) | 311 |
| Motor | Left and right hand, left and right foot, tongue | 2 (LR/RL) | 311 |
| Resting state | REST1 and REST2 | 4 (LR/RL each) | 305 |

**C. Internal imaging data**

| Source | Data/paradigm | Included data |
|---|---|---|
| Dense Amygdala release | Structural MRI | T1w/T2w individual templates; 0.9-mm source acquisitions, 0.5-mm display grid |
| Dense Amygdala release | Passive movie fMRI | *Grand Budapest Hotel* and *Forrest Gump* segments and repeats; 29 analyzed runs (372 min; 6.20 h) per participant; 2-mm slab, TR 0.556 s |
| Dense Amygdala follow-up | Emotion task fMRI | Face/shape matching; 4 runs per participant; 2-mm slab, TR 0.556 s, flip angle 48° |
| Dense Amygdala follow-up | 3D GRE TOF angiography | Two whole-brain, noncontrast arteriograms per participant; $0.26 \times 0.26 \times 0.5$ mm; registered and averaged on a 0.5-mm subject-template grid |
| Dense Amygdala follow-up | Multi-echo GRE/QSM | One 8-echo whole-brain MEGRE series per participant; $0.8125 \times 0.8125 \times 0.75$ mm; QSM-derived Frangi vesselness |

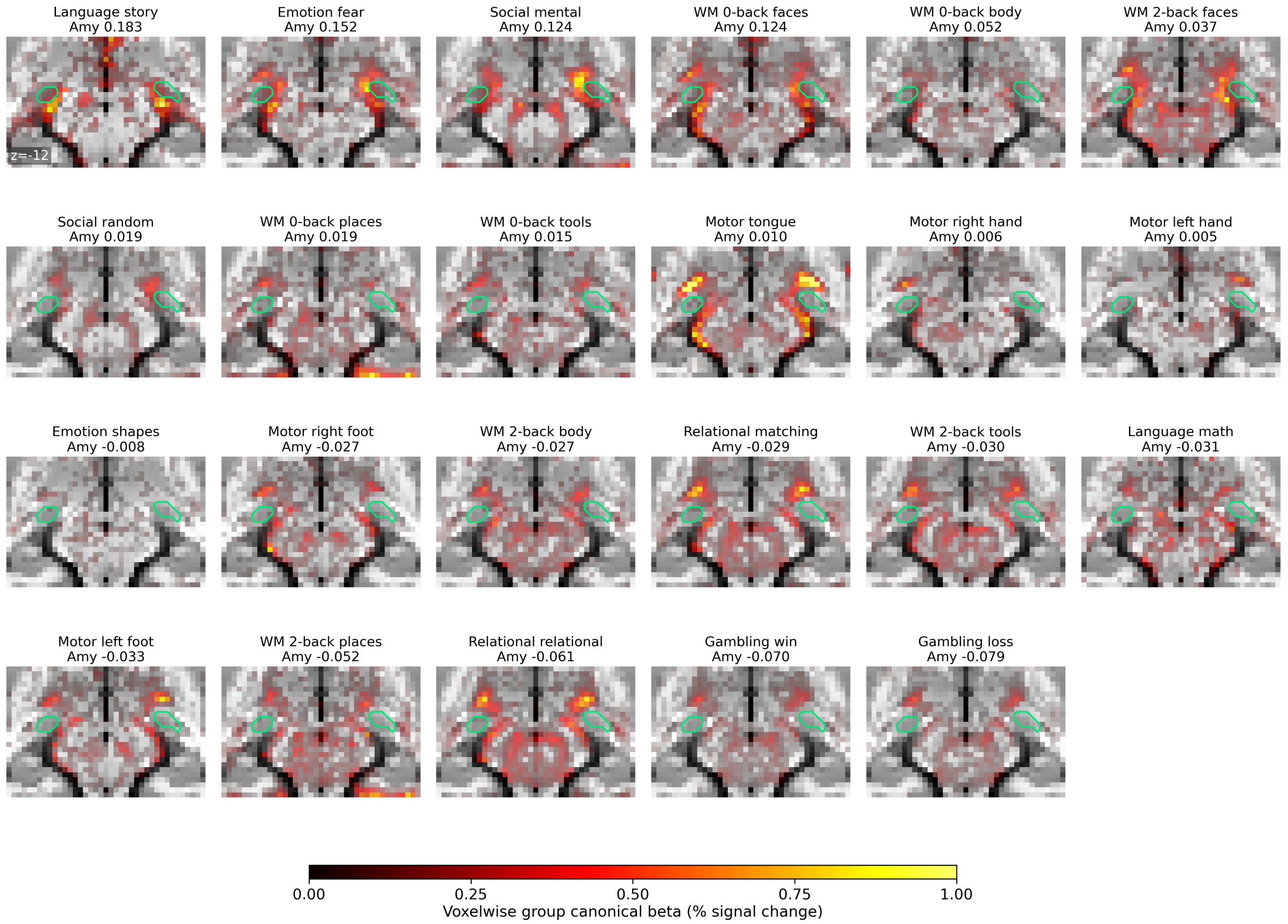


**Figure S1. Anatomical-amygdala and peri-amygdalar responses across all HCP-YA task conditions.** Group canonical-beta maps for all 23 HCP-YA task conditions, ordered by descending anatomical-amygdala response. Each panel title lists the condition and its group mean canonical beta within the CIT168 amygdala mask (Amy), in percent-signal-change units. The green contour denotes the CIT168 anatomical amygdala mask.

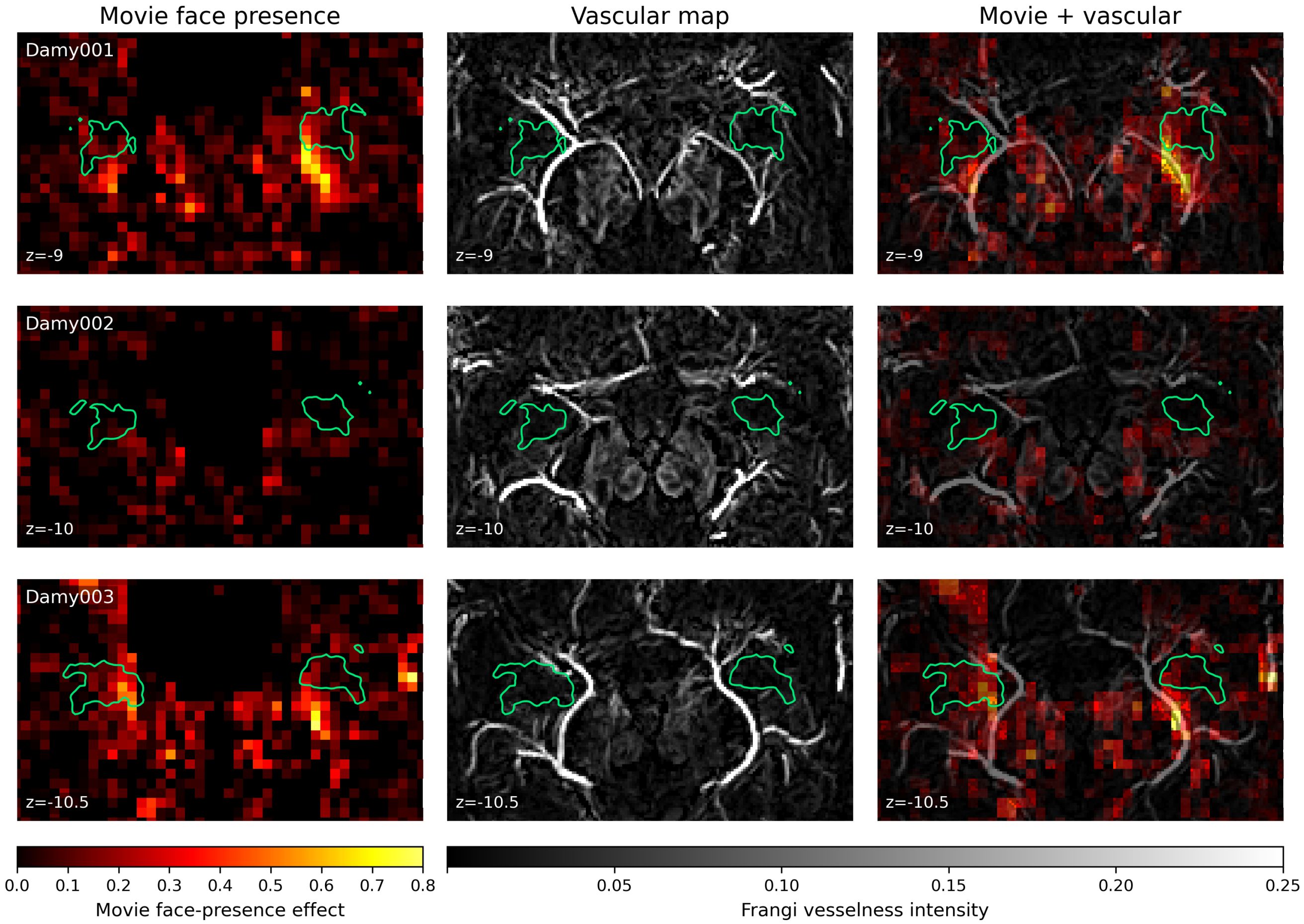


**Figure S2. Naturalistic face-presence responses reproduce peri-amygdalar overlap with subject-specific venous anatomy.** For each internal participant, the left column shows the fixed-effects face-presence effect map from the movie GLM fit to ICA-FIX-cleaned magnitude data from the Budapest and Forrest Gump movies. The middle column shows the subject-specific Frangi vesselness map rendered as a 5-mm maximum-intensity projection. The right column overlays the movie face-presence effect map and the vascular map. Green contours denote subject-specific anatomical amygdala masks. Displays use the same subject-specific axial coordinates as the internal-participant panels in Figure 3 of the main text.

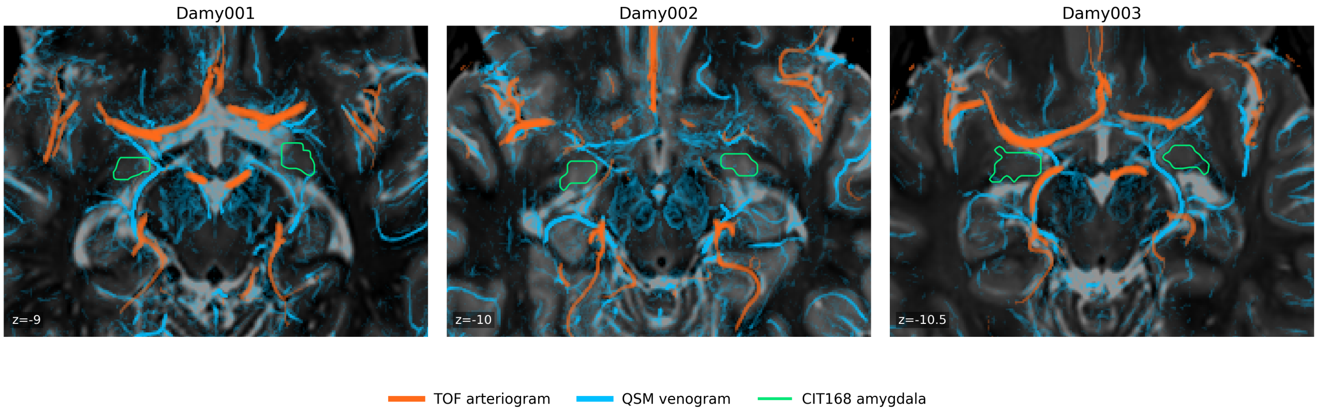


**Figure S3. Subject-specific arterial and venous anatomy in the Dense Amygdala dataset.** Columns show Damy001–003 at the subject-specific axial coordinates used in Figure 3 of the main text. Each view combines the T2-weighted anatomical template (gray), an 8-mm maximum-intensity projection of the registered and averaged TOF arteriograms (orange), an 8-mm maximum-intensity projection of the gamma-corrected QSM-derived Frangi venogram (cyan), and the CIT168 amygdala contour at $p_{\text{Amy}} \geq 0.50$ (green). All modalities were registered to each participant's 0.5-mm subject-specific template grid. These overlays are shown for anatomical context only and did not define any analysis mask.

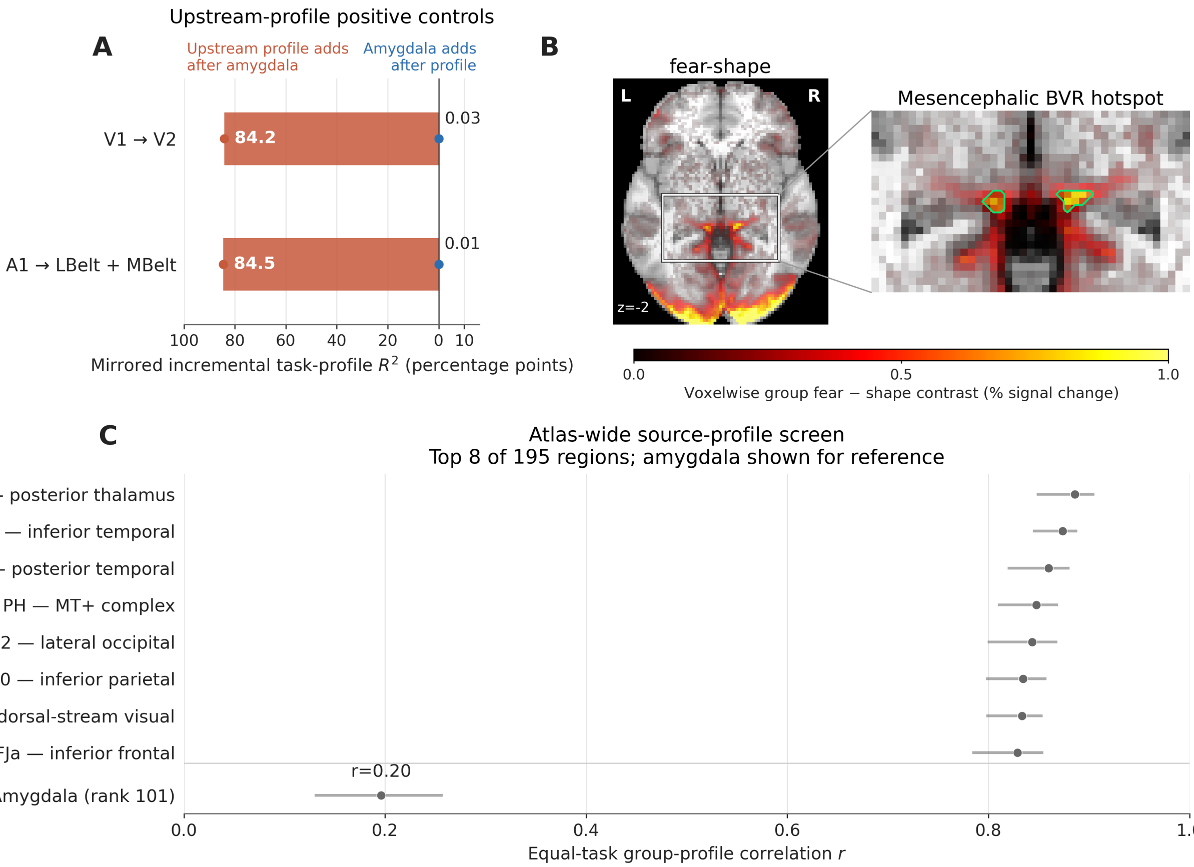


**Figure S4. Control analyses recover the expected profiles. (A)** To test whether the task-profile framework recovered expected upstream-system profiles, we applied it to two canonical controls: a V2 target with V1 as the expected upstream profile, and an LBelt–MBelt auditory-belt target with A1 as the expected upstream profile. Orange bars show variance uniquely explained by the expected profile; blue bars show variance uniquely explained by amygdala. **(B)** HCP-YA Emotion fear-shape group contrast at $z = -2$ mm. Green outlines mark the full 44-voxel mesencephalic mask. **(C)** Equal-task correlations between the 33-voxel portion of the mesencephalic mask outside the Tian S2 posterior-dorsal thalamus (THA-DP) parcel and profiles from 195 atlas-defined cortical and subcortical regions. The eight most highly correlated regions are shown, with amygdala included for reference. Points indicate full-cohort correlations, and whiskers indicate participant-bootstrap 95% confidence intervals; all 23 HCP-YA task conditions were included. The strong posterior-thalamic correspondence agrees with anatomical descriptions of posterior thalamic or pulvinar veins entering the posterior BVR. The posterior BVR also receives posterior longitudinal hippocampal and medial temporal or occipitotemporal cortical tributaries [1, 2], but these profile correspondences cannot establish parcel-specific drainage.

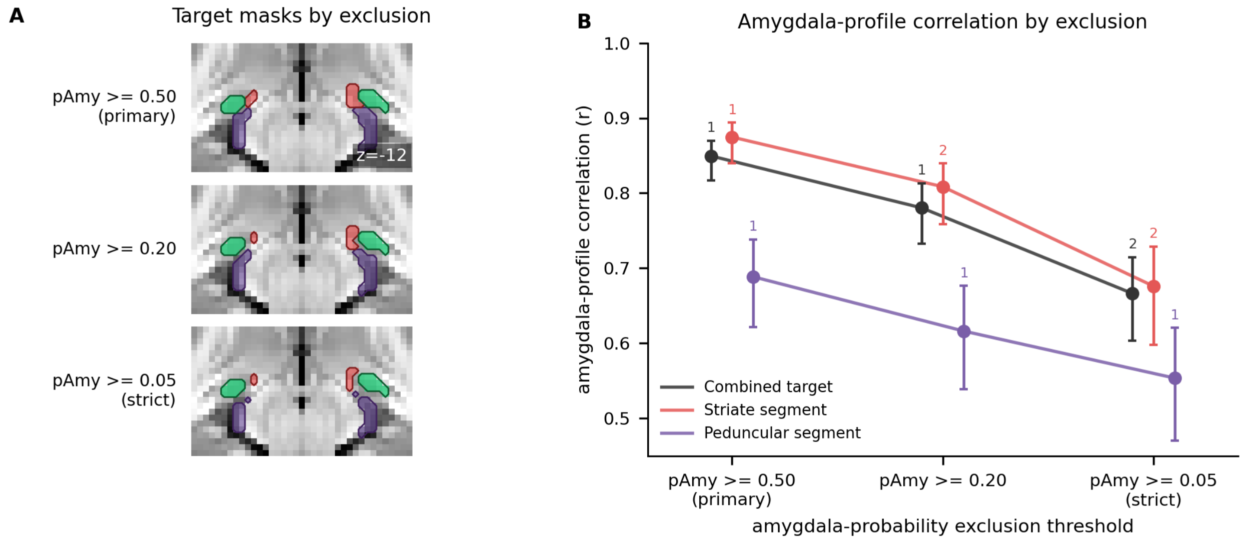


**Figure S5. Amygdala-profile correspondence weakens under stricter overlap exclusion but remains among the strongest candidate matches.** **(A)** Surviving striate (red) and peduncular (purple) target masks at each amygdala-overlap exclusion, with the CIT168 amygdala in green, on the representative axial slice ($z = -12$ mm). **(B)** Amygdala-profile correlation for the combined peri-amygdalar target and for its striate and peduncular segments after excluding voxels with CIT168 amygdala probability $p_{\text{Amy}} \geq 0.50$, $\geq 0.20$, or $\geq 0.05$. Points show full-cohort correlations, whiskers participant-bootstrap 95% confidence intervals, and labels the amygdala's rank among the 195 atlas regions. For the combined target, the amygdala ranked first in 100%, 75%, and 16% of 1,000 paired bootstrap draws at the three thresholds, respectively; piriform cortex was the leading alternative under stricter exclusions. The amygdala retained the top full-cohort rank for the peduncular segment at every threshold.

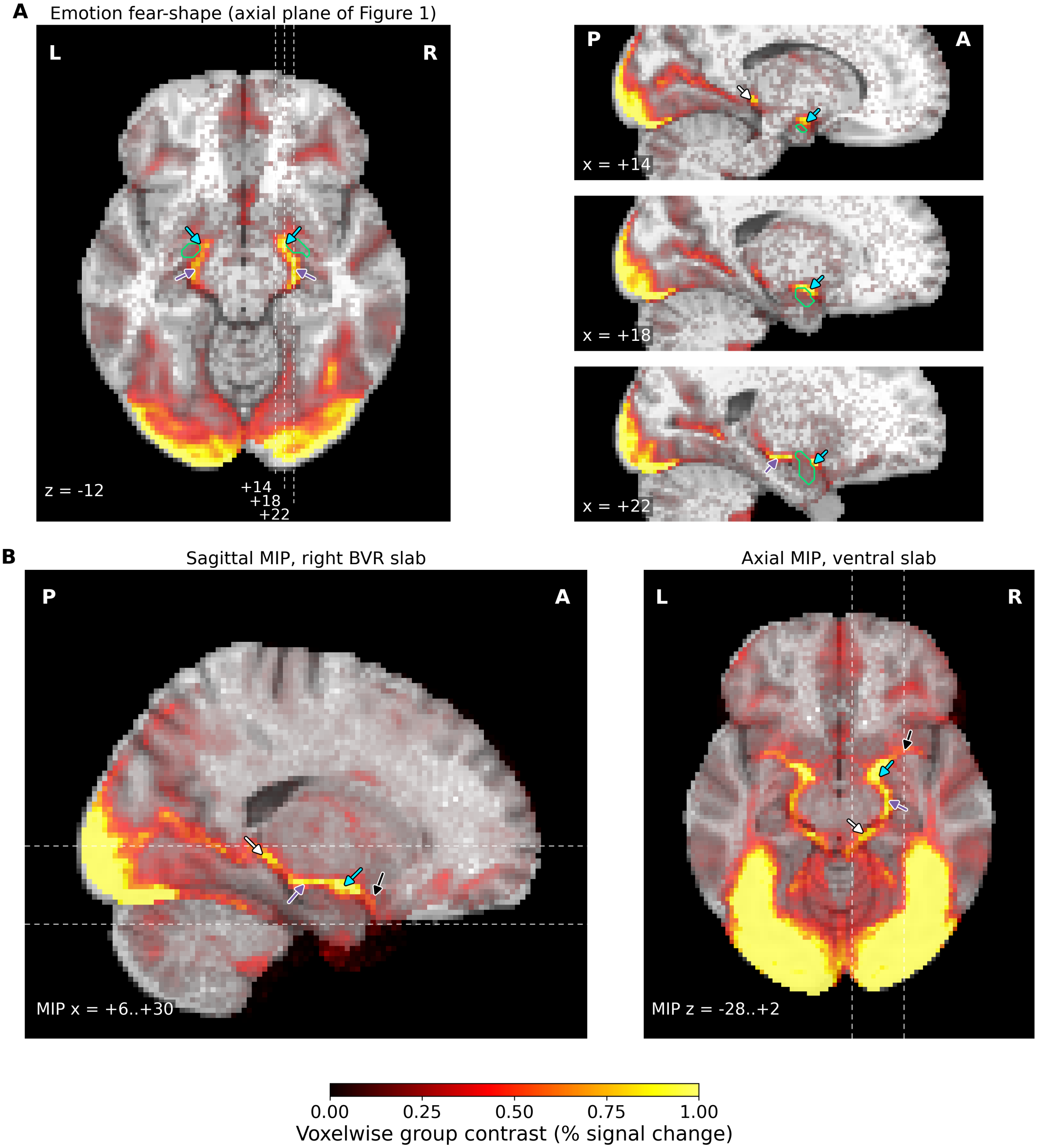


**Figure S6. Expanded-field slices and slab projections of the peri-amygdalar fear-shape response. (A)** Group emotion fear-shape contrast, using the group-map and display conventions of Figure 1 of the main text, on the same axial slice ($z = -12$ mm) and on three right-hemisphere sagittal slices ($x = +14$, $+18$, and $+22$ mm), cropped to the ventral brain. Colored arrows mark locations within the striate (cyan), peduncular (purple), and mesencephalic (white) BVR segment masks. The green contour shows the CIT168 amygdala ($p_{\mathrm{Amy}} \geq 0.50$). A similar pattern is visible in the left hemisphere. **(B)** The same contrast shown as limited maximum-intensity projections (MIPs): a sagittal MIP over the right BVR slab ($x = +6$ to $+30$ mm) and an axial MIP over the ventral slab ($z = -28$ to $+2$ mm); dashed lines in each panel mark the other panel's slab extent. Black arrows mark additional task-correlated signal anterior to the striate segment.

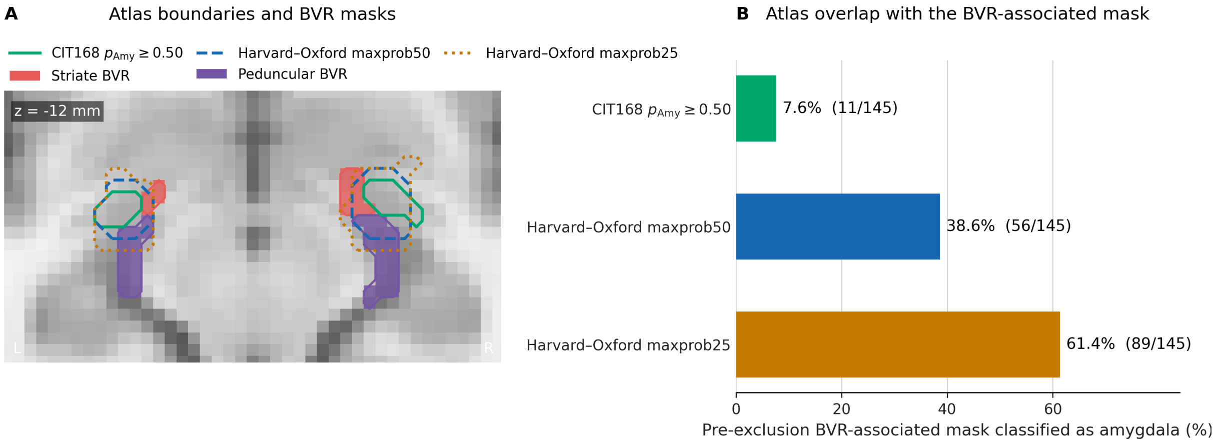


**Figure S7. Broader amygdala atlas definitions encompass more of the peri-amygdalar venous signal. (A)** CIT168 amygdala probability $p_{\mathrm{Amy}} \geq 0.50$ (green solid outline), Harvard–Oxford maxprob50 amygdala (blue dashed outline), and Harvard–Oxford maxprob25 amygdala (amber dotted outline) on a representative axial slice. The primary striate (red) and peduncular (purple) BVR masks used in the main analyses are shown after excluding voxels with CIT168 amygdala probability $p_{\mathrm{Amy}} \geq 0.50$. Harvard–Oxford maxprob50 nevertheless included 45 of their 134 voxels (33.6%). **(B)** Fraction of the 145-voxel functionally defined striate–peduncular BVR mask before atlas-overlap exclusion that was classified as amygdala by each atlas definition. At nominal 50% thresholds, CIT168 included 11 voxels (7.6%) and Harvard–Oxford included 56 (38.6%). Harvard–Oxford maxprob25 included 89 voxels (61.4%) and is shown as a threshold sensitivity analysis. This group-space comparison demonstrates sensitivity to atlas boundaries; neither atlas is an individual-level histological or venographic ground truth.
